# A multiplexed work-flow for absolute quantification of *Klebsiella oxytoca* Nif proteins

**DOI:** 10.1101/2020.04.27.064014

**Authors:** Christoph James Waite, Anya Lindstrom Battle, Mark Bennett, Martin Buck, Jörg Schumacher

## Abstract

Global imbalances of the nitrogen cycle are increasingly recognized as a major challenge to human development, and are exacerbated by the use of synthetic inorganic fertilizers. Biotechnology alternatives to inorganic fertilizers include biofertilisation from nitrogen fixing bacteria (diazotrophs), expressing nitrogenase and auxiliary genes (*nif* genes). In order to directly quantify all twenty Nif proteins through multiple reaction monitoring (MRM) MS, we established a high throughput pipeline to generate a set of *Klebsiella oxytoca* Nif protein QconCATs as quantotypic standards. A stringent validation of the pipeline and QconCATs application with regards to isotopic labelling efficiency (100%), limits of detection and quantification, analyte to internal standard concentration boundaries was used for optimisation. Using three QconCATs for the measurement of 20 likely low, middle and high Nif protein abundances enabled detection of all Nif proteins, 19 of which could be accurately quantified and their variation over time monitored. Stoichiometries between *Klebsiella oxytoca* Nif proteins and changes between early and late transition into diazotrophy suggest i) a temporal regulation of *nif* gene cluster expression that may be linked to nitrogenase expression and maturation; ii) vast disparities in Nif protein abundances and iii) high dependency on the *nifLA* master regulator pair for *nif* gene expression.

## Introduction

Global imbalances of the N cycle are increasingly recognized as a major challenge to human development (1). Nitrogen availability is a major limiting factor for crop growth, such that nitrogen cycles drive agricultural productivity. Specialised bacteria and archaea, known as diazotrophs, have evolved the capability to fix atmospheric nitrogen to NH_3_ before assimilating it for use in biological processes, using a series of nitrogen fixation (*nif*) genes (2). Biological nitrogen fixation is an important part of the nitrogen cycle, as it replenishes the nitrogen content of the biosphere. Nitrogen fixation is accomplished by the highly conserved Adenosine triphosphate (ATP)-hydrolysing and redox-active nitrogenase enzyme complex(3). *Klebsiella oxytoca* (previously classed as *K. pneumonia*), a free-living diazotroph, has long been used as a model system for the study of N fixation. Nitrogenases are composed of a reductase (Fe protein) and a catalytic component (MoFe protein). The MoFe protein is a tetramer of NifD and NifK, carrying an iron-molybdenum cofactor (FeMo-co[MoFe_7_S_9_-C-homocitrate]) in each NifD subunit and a P-cluster ([8Fe-7S] cluster) at each NifD-NifK interface. NifH forms the dimeric Fe protein, with a [4Fe-4S] cluster in the subunit interface (4). Most *nif* gene products are involved in the assembly, function and regulation of the biochemically complex nitrogenase. In *K. oxytoca, nif* genes organised into 7 operons, enabling co-ordinated expression (5). The assembly of active nitrogenase requires the Nif proteins to function together either in series or in tandem. The NifU/NifS machinery is sufficient to provide [4Fe-4S] for NifH biogenesis. NifDH, however, requires many Nif proteins for biosynthesis of the structurally complex FeMo-co. NifU/NifS generate the NifB cofactor by providing [Fe-S] cluster substrates, which are transferred to NifEN. Fe, Mo, and homocitrate are incorporated into FeMo-co by NifQ, NifH, NifV, NifT and NifX. The completed FeMo-co is incorporated into apo-NifDK via NifY to yield holo-NifDK (8). An overview of *nif* genes and their functions in *Klebsiella oxytoca* is provided in table 1.

**Table 1:**
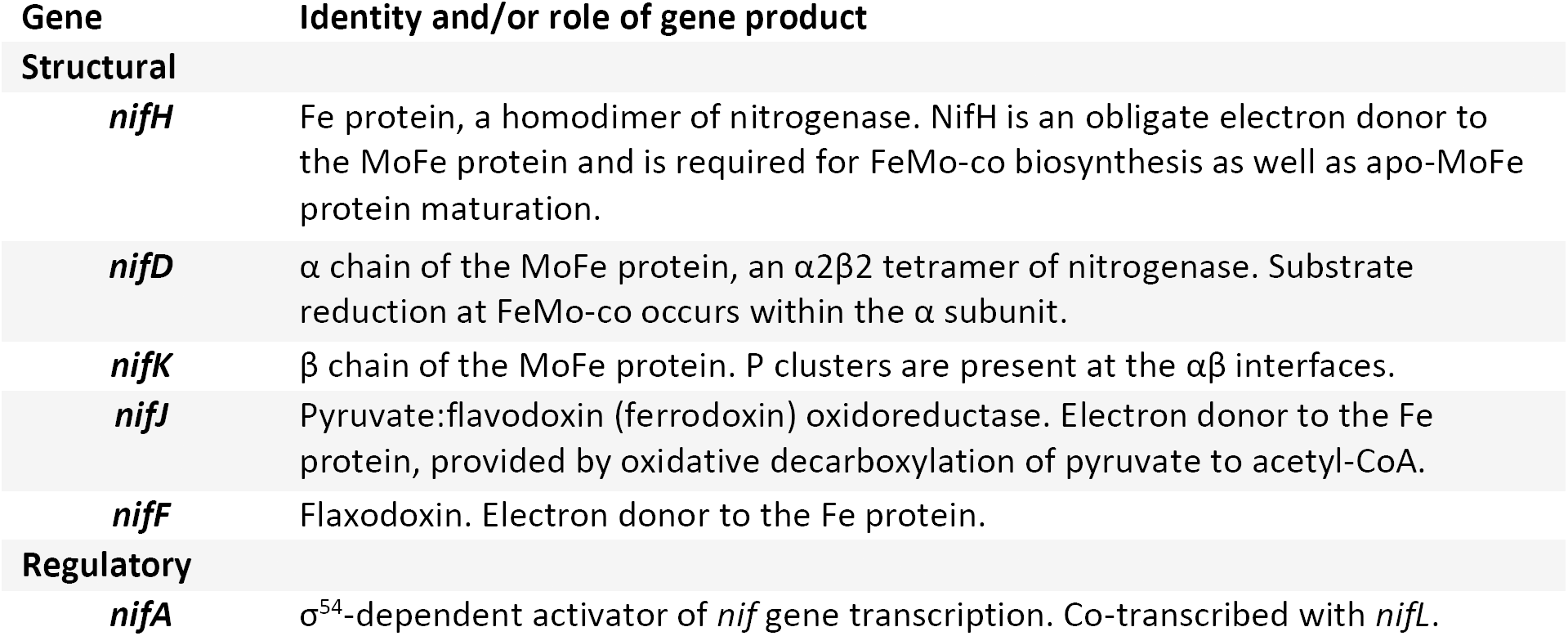
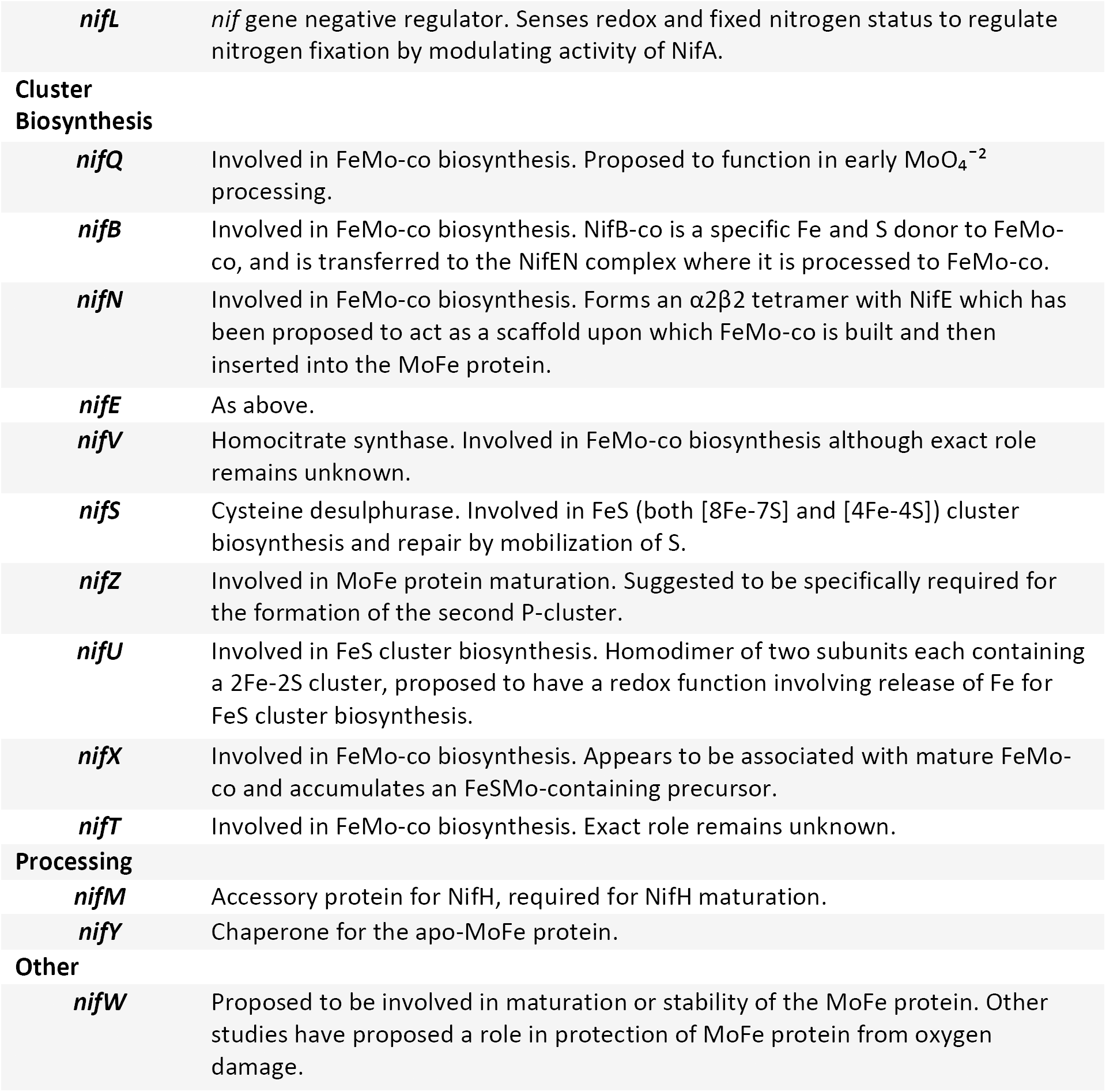
*Nif* genes *of Klebsiella oxytoca* M5a1 and their functions.

In diazotrophic proteobacteria, such as *K. oxytoca*, the two-component system NtrB-NtrC couples N limitation to a global transcription response upon NtrC phosphorylation (13). The level of phosphorylated NtrC controls the σ^54^-dependent transcription of many genes associated with nitrogen assimilation, in addition to those encoding key regulators of *nif* gene cluster expression, *nifL* and *nifA*. NifA initiates transcription of the remaining six *nif* operons via σ^54^-dependent promoters. However, NifA activity is repressed upon interaction with NifL in conditions unsuitable for nitrogen fixation, such as oxygen and increased fixed N (6). N status regulates *nif* gene expression at both the transcriptional (via NtrC) and post-translational (via NifL) levels. Post-translational modifications play a key role in rapidly fine-tuning the N fixation and assimilation pathways in response to internal N status, specifically glutamine concentration (14). The PII-like proteins GlnK and GlnB are covalently modified by the uridylytransferase (UTase) GlnD in response to low glutamine. In contrast, glutamine stimulates the uridylyl-removing (UR) activity of GlnD. By modulating the NifA-NifL interaction, GlnK is responsible for additional N-dependent regulation of *nif* gene expression (9). The NifL-NifA interaction is inhibited merely by the presence of GlnK, irrespective of uridylation status. However, covalent modification of GlnK becomes important in its interaction with the membrane-associated ammonium transporter AmtB. The de-uridylation of GlnK in response to increased nitrogen allows for membrane sequestration of GlnK by AmtB, decreasing its cytoplasmic concentration for modulation of NifL-NifA interaction (16). The other PII-like protein GlnB modulates the kinase/phosphatase activities of NtrB upon direct sensing of glutamine, but only when de-uridylated (6).

Advances in multiple reaction monitoring mass spectroscopy MRM-MS are applied to quantify targeted proteins in complex biological samples with high sensitivity. Triple quadrupole mass spectrometers are best suited for protein absolute standard quantification, where the three chambers (Q1, Q2, Q3) can be set to act as mass filter, collision chamber and fragment ion detector and selected precursor-fragment ion pairs (termed transitions) can be measured directly. Quantitative proteomics commonly involve the addition of labelled standards to biological samples, against which the protein of interests can be measured in the mass spectrometer. MRM-MS is both extremely sensitive, detecting peptides down to the attomole range, and extremely selective. While label-free quantitative approaches can measure hundreds of proteins simultaneously, the availability of labelled standards has several advantages with regards to accuracy, precision, reproducibility and stoichiometry (reviewed 6).

The addition of isotopically labelled standards of known concentration to the sample of interest allows the absolute quantification of targeted sample proteins in MRM-MS. Various isotopically labelled peptidic standard approaches have been used for absolute protein quantification: i) protein standard absolute quantification (PSAQ), where the protein to be measured is purified and isotopically labelled (7); ii) AQUA (absolute quantification) involves the addition of labelled peptides to be measured and QconCATs are artificial proteins encoding a concatemer of tryptic peptides specific to a protein set of interest in a single open reading frame facilitates multiplexed protein quantification (24, 25). The choice of which labelled standard approach is commonly guided by how many proteins are targeted (6). In the 10-50 target protein range, QconCAT fares best, and was adopted here to quantify the 20 Nif proteins. These peptides must be ‘quantotypic’, in that they are unique to the protein under study and suitable for quantification (26). Endoproteolytic fragmentation of the QconCAT-analyte mix results in release of QconCAT peptides in a strict 1:1 ratio, which can serve as internal control during analysis. To further improved the QconCAT methodology, we increased trypsin digestion reproducibility between target and standard proteins by introducing target amino acids N-terminal to the trypsin cleavage site. For the determination of intracellular protein concentrations we optimised sample preparation and measurements in order to increase reproducibility across biological samples. For the results shown in table two of the main document, standard errors of the mean across three biological replicates ranged from 0.8% to 13%, demonstrating high robustness of the workflow.

## Materials and Methods

QconCAT design: *In silico* trypsin digestion of annotated *K. oxytoca* M5a1 Nif protein sequences, obtained from NCBI accession code AMPJ00000000, and peptide uniqueness analysis were undertaken in Skyline (10). For identification of quantotypic peptide candidates, the Skyline-generated transition list was exported and run against Nif protein-containing *K. oxytoca* M5a1 protein extract prepared as outlined below. Final peptide selection was based on the following favourable characteristics: absence of Cys and Met, absence of His, presence of Pro, Pro not adjacent to cleavage site, lack of residues with possible artefactual modification (Glu, Trp, Asn, N-terminal Gln, adjacent Lys/Arg/His at either terminus), favourable peak characteristics (multiple co-eluting transitions, symmetrical peak (11,12,13). Lastly, peptides were chosen by retention time to maximize peak separation. Initial control for peptide identity was done by comparing against a control *K. oxytoca* M5a1 protein extract expected to not contain any Nif proteins (aerobic, Δ*nifLA*). An Enhanced Product Ion (EPI) scan was used to confirm peptide identity. Known tryptic peptides from Bovine Serum Albumin (BSA) identified from previous Multiple Reaction Monitoring MS (MRM-MS) experiments were selected based on retention time and placed on the 3’ and 5’ end of each QconCAT, to act as internal controls against potential degradation by exoproteolysis (13). Each QconCAT was commercially synthesized (GeneArt, Life Technologies). A final transition list was exported from Skyline and used for all subsequent analyses (Appendix I).

### Growth of K. oxytoca under anaerobic conditions

Single colonies of wild-type (WT) *K. oxytoca* M5a1 and *K. oxytoca* M5a1 Δ*nifLA* were used to inoculate a 2 ml preculture in Luria-Bertani (LB) media (5 g/l NaCl, 5 g/l yeast extract, 10 g/l tryptone). Cells were grown at 30 °C at 200 RPM until reaching a suitable density, and 10 ml overnight cultures set up in NFDM media (2 ml 5X NFDM [100 g/l glucose, 2 mM MgSO_4_, 0.5 mM Na_2_MoO_4_, 0.45 mM FeSO_4_, 125 mM KH_2_PO_4_, 350 mM K_2_HPO_4_], 200 μl 1 M NH_4_Cl) with 25 μl WT or 50 μl Δ*nifLA* cell suspension, and grown at 30 °C overnight.

Cultures were centrifuged at 4000 g and pellet washed with 1X NFDM to remove residual ammonia. Cells were resuspended in 10 ml 1X NFDM and kept on ice to temporarily inhibit cell growth. The OD_600_ of re-suspended cultures was measured using a SpectroStar Nano microplate reader (BMG LABTECH) and normalised volumes of each culture were added to 40 ml cold NFDM media (as above, scaled up) in sealable glass culture vessels for a final OD_600_ of 0.1. WT *K. oxytoca* M5a1 was grown in both 10 (N-rich) and 0.25 (N-poor) mM NH_4_Cl, and Δ*nifLA* (mutant) in 0.25 mM NH_4_Cl. Vessels were fixed with airtight caps and sparged with filter-sterilised N_2_ for at least 25 minutes. One vessel was checked for absence of oxygen using O2xyDot® technology (OxySense). Finally, 1 ml of pure acetylene was added before the vessels were warned to 25 °C. OD_600_ and NH_4_ concentration were measured regularly in all cultures by sampling 0.4 ml using a N-sparged 1 ml syringe. OD_600_ was measured as described above following a 1/4 dilution of culture. Supernatant for NH_4_ assay was obtained by centrifugation, as above, of 150 μl culture and storing at 4 °C. Acetylene reduction by gas chromatography-MS (GC-MS) was checked at t = 4, 8.5 and 10 hours (Appendix IV). Early and late N-fixing transition samples were taken according to results, at t = 5 and t = 10. 15 ml of the cell suspension was sampled and centrifuged for 25 minutes at 4000 g before cell pellets were frozen at -80 °C. [NH_4_] was assayed the next day using the Ammonium Test (Merck Millipore) and a standard curve (R^2^ = 0.99 [to 2 decimal places]) of duplicates of 10 to 0 mM NH_4_ in culture media.

### QconCAT expression and purification

Designed QconCATs were received from Life Technologies in a pET100/D-TOPO vector with a 5’ hexa-histidine tag, an ampicillin resistance gene, and a *lacO* operator. Transformation of CaCl_2_-chemically competent *E. coli* BL21 Δ*lys,arg* cells was undertaken according to standard protocols before plating on LB agar 100 μg/ml ampicillin-selection plates. Each QconCAT was expressed in ^13^C_6_^15^N_4_ arginine and ^13^C_6_^15^N_2_ lysine-labelled expression media (1X Gutnick media [33.79 mM KH_2_PO_4_, 77.51 mM K_2_HPO_4_, 5.74 mM K_2_SO_4_, 0.41 mM MgSO_4_·7H_2_O], 0.4 % glucose, 10 mM NH_4_Cl, 1 mM of each amino acid, 1X HoLe trace elements) (Appendix II). Overnight cultures were inoculated by adding one colony from each QconCAT plate to 10 ml labelled media with 100 μl/ml ampicillin, and were incubated with shaking at 37 °C. 2 ml of the overnight culture was added to 100 ml pre-warmed labelled media. Day cultures were grown to an OD_600_ of 0.4-0.6 at 37 °C with shaking, at which point 1 ml uninduced cell sample was removed. Cells were induced by adding isopropyl β-D-1-thiogalactopyranoside (IPTG) to a final concentration of 1 mM and incubated for 4 hours at 37 °C with shaking. 1 ml of each induced culture was removed, and cultures were centrifuged for 20 minutes at 4000 g before storing at -80 °C. Induced and uninduced cell samples were centrifuged at 4000 g for 5 minutes before storing pellet at -80 °C.

The pellet was resuspended in 20 ml 20 mM Tris-HCl (pH 8.0) before pulsed sonication (2 seconds on, 2 seconds off, 40% amplitude) for 1 hour on ice. The broken cell suspension was centrifuged at 4 °C for 45 minutes at 27000 g, 8 μl of the supernatant aliquoted for sodium dodecyl sulphate polyacrylamide gel electrophoresis (SDS-PAGE) and stored at -80 °C. The pellet was resuspended in 20 ml 20 mM Tris-HCl with 7 M urea. The suspension was centrifuged as before and supernatant aliquoted.

4 ml new Ni-NTA resin per purification was prepared by a series of centrifugation (4 °C, 4000 g, 5 minutes) and wash (4 ml water, 4 ml binding buffer [95 mM Na_2_HPO_4_, 5.2 mM NaH_2_PO_4_, 50 mM NaCl, 3 M urea]) steps. 9 ml of the QconCAT-containing supernatant was added to the resin and incubated at room temperature for 45 minutes with gentle shaking. The mix was added to a purification column and flow-through collected. Buffers were added sequentially and flow-through collected: 25 ml binding buffer, 15 ml wash buffer (binding buffer + 30 mM Imidazole), 2 ml elution buffer (binding buffer + 500 mM Imidazole). 8 μl aliquots of each flow-through were collected for SDS-PAGE. The elution was dialysed in 12-15 kDa molecular weight cut off dialysis tubing in dialysis buffer (1X Gutnick, 7 M urea) overnight at 4 °C with stirring. Dialysis buffer was replaced and sample dialysed for at least 1 additional hour. 1 mM tris(2-chloroethyl) phosphate (TCEP) was added and the purified QconCAT stored at -80 °C.

### SDS-PAGE

15 % acrylamide SDS-PAGE gels were prepared and run according to the Laemmli method. 4 μl 3X Instant-Bands Sample Treatment Buffer (EZ Biolab) was added to 8 μl sample before incubation at 95 °C for 5-10 minutes. 10 μl of samples was run against 3 μl of the EZ-Ladder Fluorescent Protein Molecular Weight Markers (EZ Biolab) at 200 V for 60 minutes. The gel was visualized under ultraviolet (UV) light using a Gel Doc 2000 (BIO RAD). For the induced and uninduced cell samples, 100 μl 2X Laemmli buffer (4 % SDS, 20 % glycerol, 120 mM Tris-HCl [pH 6.8], 0.02 % bromophenol blue) and 1 μl pure β-mercaptoethanol were added, before incubation at 95 °C for 5 minutes. Samples were centrifuged for 10 minutes at 14000 g, and 10 μl run against 5 μl PageRuler Prestained Protein Ladder (ThermoFisher Scientific) as above. Gel was stained with Coomassie-based SimplyBlue SafeStain (ThermoFisher Scientific). Concentration of the QconCATs was established by a detergent compatible (DC) Assay (BioRad) according to the manufacturers protocol. A standard curve (R^2^ = 1.00) with 9 concentrations ranging from 0 to 2 mg/ml BSA in duplicates was used.

### MRM-MS sample preparation

The sample pellets were resuspended in 1 ml 1X Gutnick with 7 M urea before sonication for 10 minutes on ice as before. Sample was centrifuged at 15000 g and 4 °C for 45 minutes. DC assays were undertaken on protein-containing supernatant as above. Trypsin digestion reactions were set up as follows: 30 μl protein extract, 3 μl of each QconCAT at appropriate dilution (see figure 7c), 3 μl 100 mM TCEP, 50 mM NH_4_CO_3_ up to total of 290 μl. An 8 μl pre-digestion sample was taken. 5 μl trypsin at 0.4 mg/μl was added and the reaction incubated at 37 °C for 4 hours, before incubation at room temperature overnight. An 8 μl post-digestion aliquot was taken and 15 μl pure formic acid (5%) added. Samples were centrifuged at 15000 g prior to pipetting into liquid chromatography (LC) vial.

### QconCAT validation

All MRM-MS results were viewed and analysed on Skyline, and quantification based on integrated peak area. Manual integration of all peaks was undertaken, such that (a) the correct peak corresponding to the tryptic peptide was selected and (b) integration boundaries defined the area between points where the curve first touches the x-axis. Manual integration was adjusted if Skyline-calculated background was >5 % of total peak area. The Δ*nifLA* and WT N-poor samples at t = 10 were used to vary the % Nif protein in the Nif protein signal linearity assay. After adjusting samples to the same total protein concentration, samples were mixed such that Nif protein-containing sample constituted 100, 50, 10, 1, 0.1, 0.01 and 0.001 % of the protein extract. Linearity of High QconCAT signal was assayed by preparing a dilution series such that QconCAT concentration added to the trypsin digestion reaction was 2, 1, 0.2, 0.04, 0.008, 0.0016 times stock concentration. Trypsin digestion was set up as above and samples run with final QconCAT transition list (Appendix I). Data analysis using Skyline was undertaken as described, and absolute quantification values obtained from standard:sample ratio and known concentration of isotope-labelled QconCAT. Quantification as copies per cell was derived from OD_600_ (supplementary file).

### Mass Spectroscopy

For quantification, detection of the compounds was based on MRM-MS. Samples (typically 40 μl) were analysed by triple quadrupole HPLC-electrospray ionisation/MS-MS using a Shimadzu Prominence UPLC coupled to an Applied Biosystems Q-TRAP 6500 (AB SCIEX, Massachussetts, USA). Chromatographic separation was carried out by reverse phase chromatography on a Phenomenex Luna C18(2) 100A (3 μm, 100 by 2 mm) HPLC column, at 50 °C. A gradient system using A (5 % CH_3_CN, 94.9 % H_2_O, 0.1 % formic acid) and B (5 % H_2_O, 94.9 % CH_3_CN, 0.1 % formic acid) was used, with solvent flow rate of 250 μl/minute. A linear gradient from 0 % B to 25 % B over 30 minutes was followed by an increase to 50 % B over the next 5 minutes. B was then increased to 100 % over the next 2 minutes and held at 100 % for 5 minutes, before return to initial conditions. The MS was operated in the positive mode using an IonDrive™ Turbo V as the ion source. Source conditions were: temperature, 500 °C; ion source gas 1, 40 psi; ion source gas 2, 60 psi; ion spray voltage, 5500 V; curtain gas, 40 psi; CAD gas setting, -2. An Enhanced Scheduled MRM method was used for the analysis, and details of the full method can be found in appendix I. For qualitative peptide validation, the MS was run in ‘trap’ mode for an Enhanced Product Ion (EPI) scan using the same source conditions. An information dependent acquisition (IDA) method was carried out based on (29). For qualitative peptide validation, the MS was run in ‘trap’ mode for an Enhanced Product Ion (EPI) scan using the same source conditions. An information dependent acquisition (IDA) method was carried out based on (14).

## Results

### Growth and N fixing profiles of N-poor, N-rich and mutant K. oxytoca strains

To establish growth conditions for *K. oxytoca* M5a1 and *K. oxytoca* M5a1 Δ*nifLA* (mutant), cells were grown in NFDM medium under anaerobic conditions over a time course of 9.5 hours, were growth and N fixing parameters were monitored hourly. The concentration of NH_4_ in the N-rich sample steadily decreased over the course of 9.5 hours. In the case of both the mutant and N-poor cultures, [NH_4_] decreases steadily until t = 2, at which point it runs out (Figure 1). Acetylene reduction was observed only in the WT N-poor culture, from at least 4 hours after N-sparging (Figure 4c). Samples at 5 and 10 hours were deemd early and later N fixation states in *K. oxytoca*.

**Figure 1:**
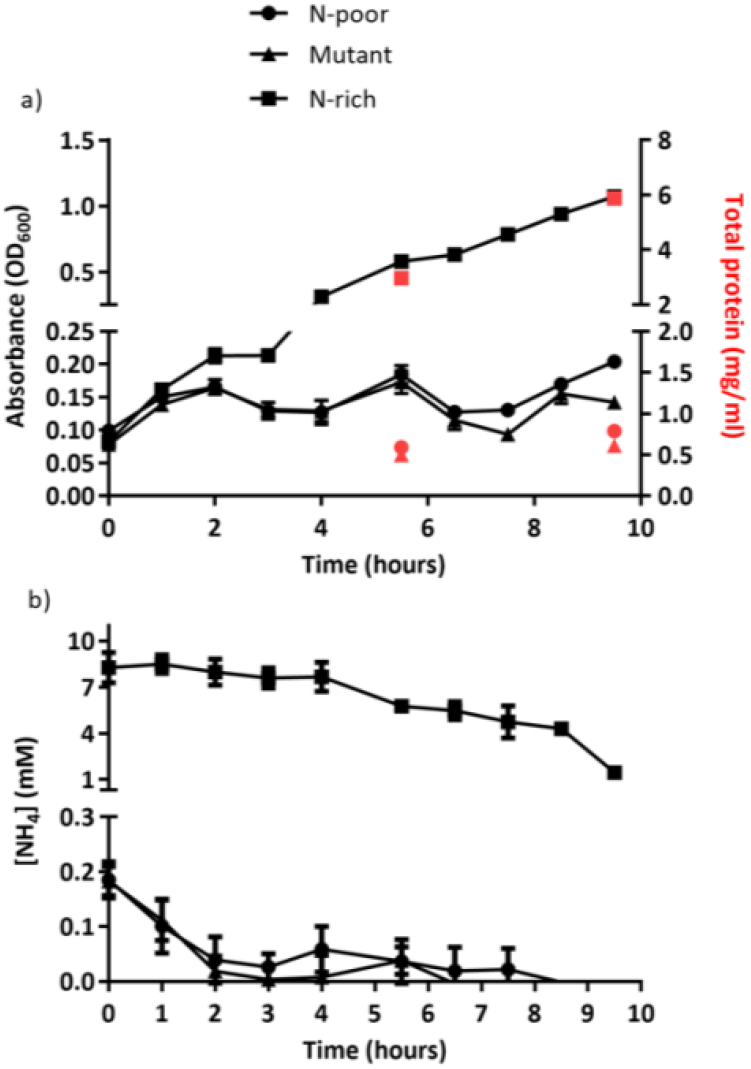
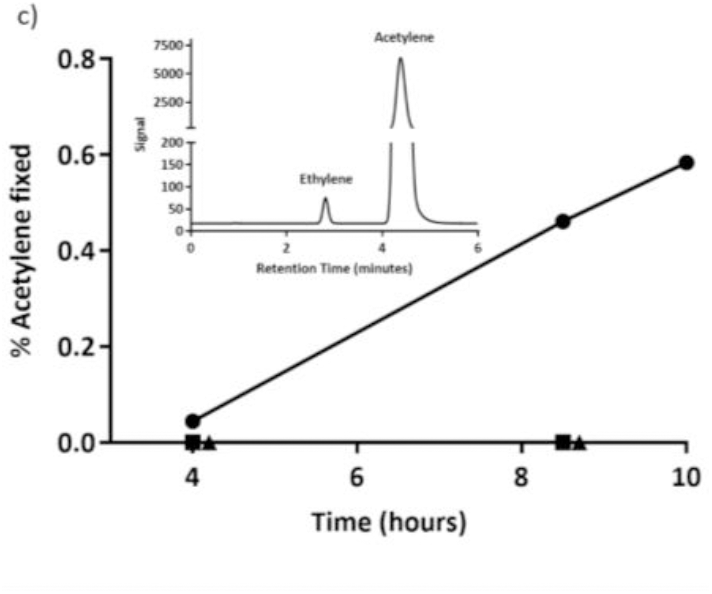
Growth and nitrogen fixation of *K. oxytoca* M5a1 grown in 0.25 (N-poor) and 10 (N-rich) mM NH_4_, and *K. oxytoca* M5a1 Δ*nifLA* (mutant) grown in 0.25 mM NH_4_, all under anaerobic conditions. (a) Growth profiles of N-poor, N-rich and mutant cultures in terms of absorbance (OD_600_) from 0 to 9.5 hours after N sparging. (b) Ammonium concentration of the supernatant of each culture was measured by the Merck Millipore Ammonium test. In (a) and (b), error bars represent the Standard Error of the Mean (SEM) over three biological replicates. (c) One replicate of each culture was tested for N fixation by conversion of acetylene to ethylene at t = 4, 8.5 and 10 hours after N sparging. % acetylene conversion was calculated by peak area changes in GC-MS results. Inset is the GC-MS result obtained at t = 10 hours for the N-poor sample showing the expected appearance of an ethylene peak at approximately 2.5 minutes. Error was not calculated for (c) as only one sample was taken at each time point. Points representing % acetylene for the mutant are offset by 0.25 for clearer presentation. Note that in (a), (b) and (c) inset the y-axis is split.

**Figure 2:**
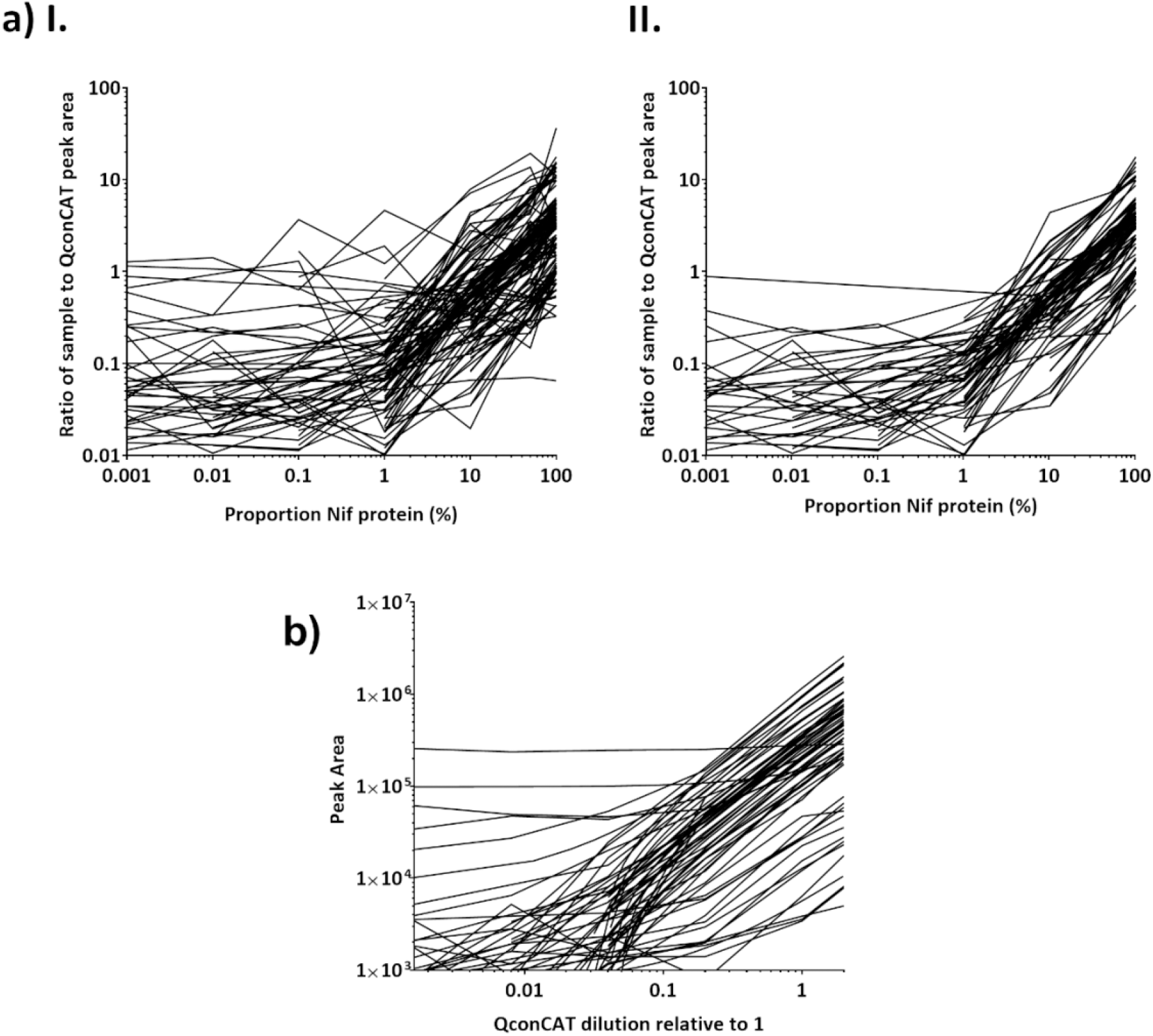
Validation of QconCAT-mediated quantification of Nif protein by MRM-MS. (a) A Nif protein-containing sample dilution series was established in order to identify transitions reliably quantifiable (linear), and over what range (I.). Transitions which were not linear were removed (II.) and were not included in further analyses. Ratio of the peak area of each analyte peptide to the corresponding heavy QconCAT peptide were graphed on a logarithmic scale against % Nif protein in the sample. (b) The High QconCAT was diluted and peak areas calculated to identify the linear, and thereby quantitative, range. All peak areas were calculated by Skyline as is outlined in Methods.

**Figure 3:**
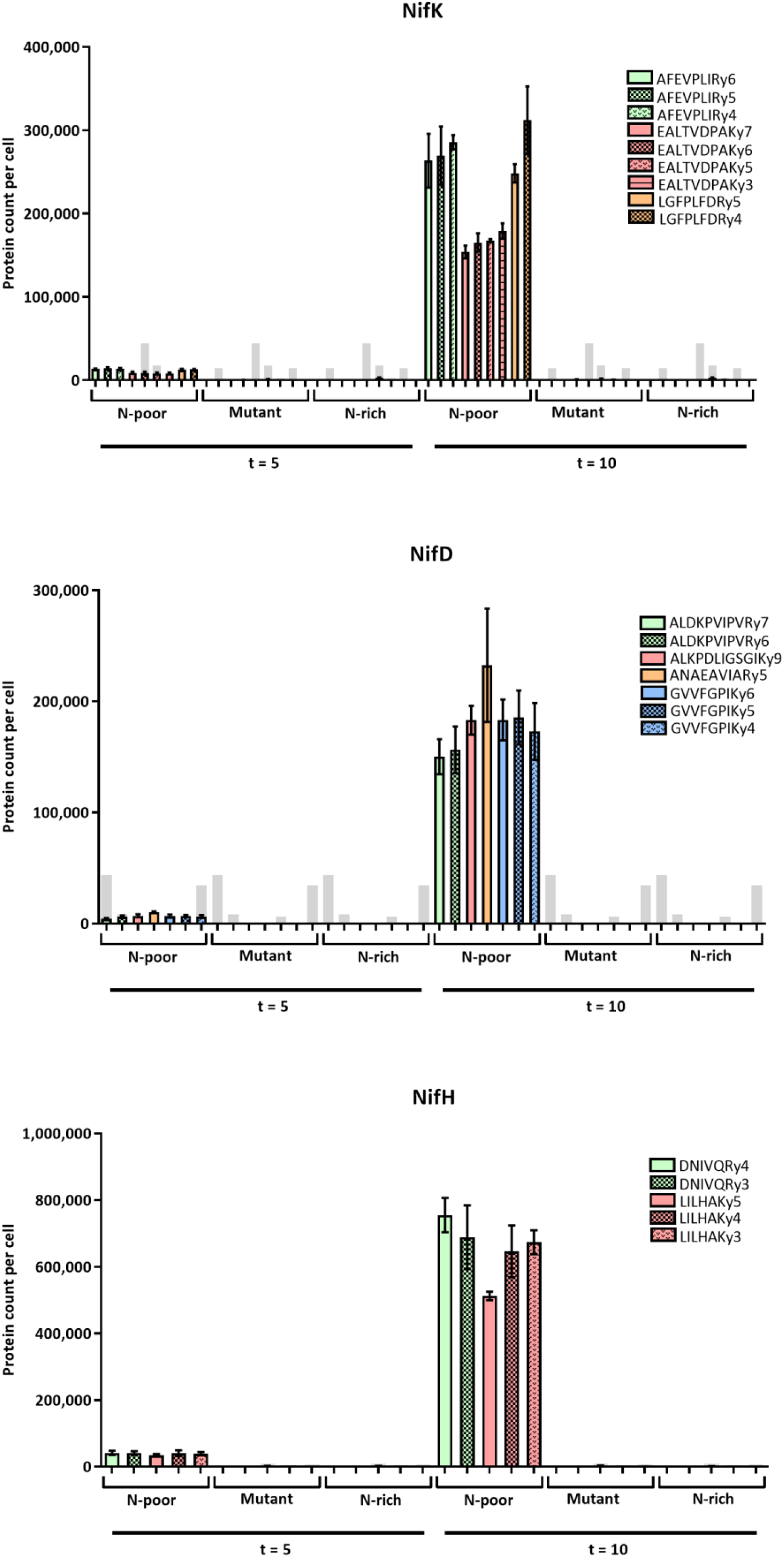
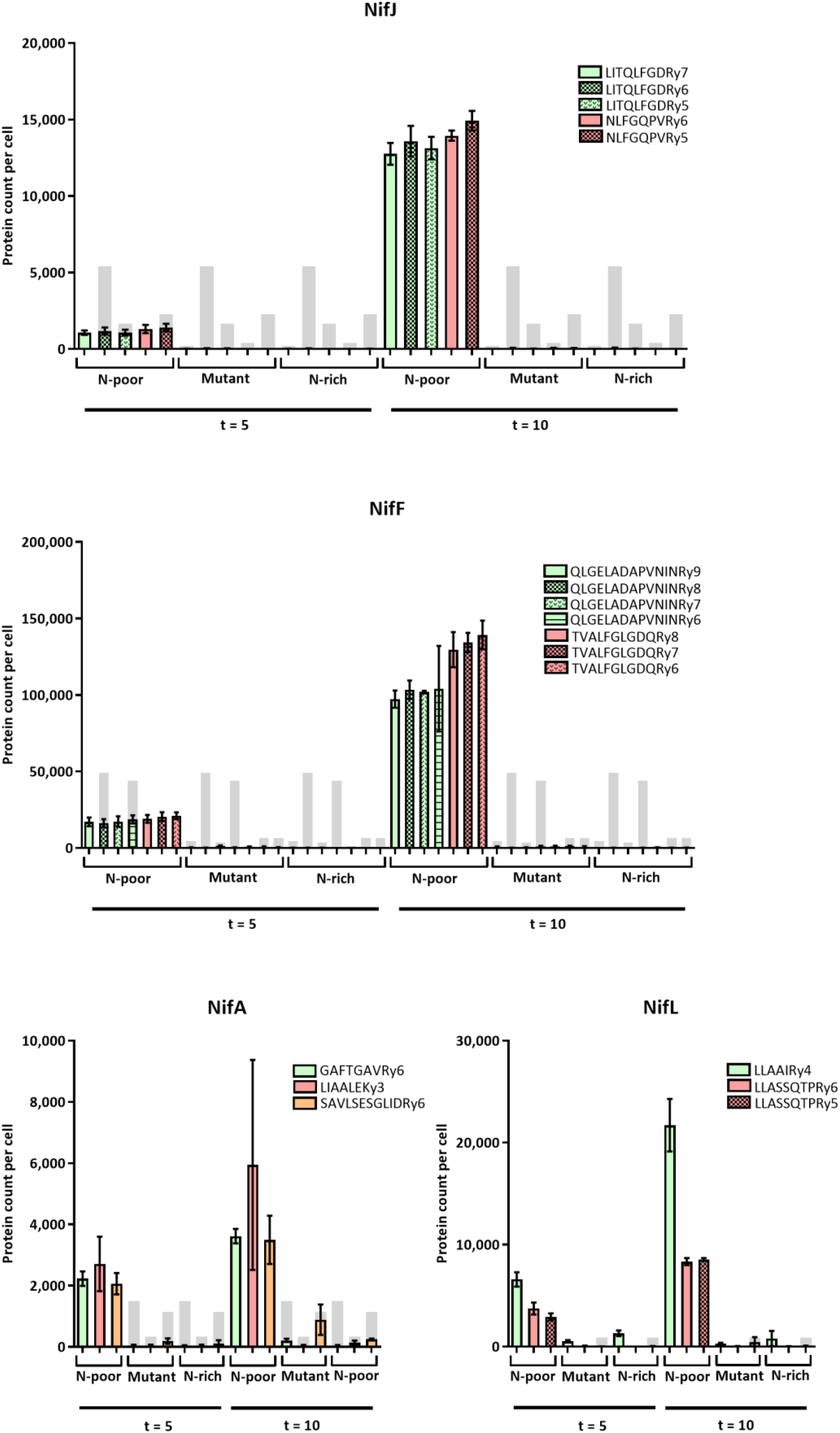
Protein count per cell for all individual Nif proteins in K. *oxytoca* M5a1 grown in 0.25 (N-poor) and 10 (N-rich) mM NH_4_, and *K. oxytoca* M5a1 Δ*nifLA* (mutant) grown in 0.25 mM NH_4_, at both 5 and 10 hours after N sparging. All transitions for each peptide are shown, and limit of quantification illustrated by a filled in light grey bar. Error bars represent SEM over three biological replicates. Protein counts for all other Nif proteins are provided in supplementary file. Note that y-axis scale varies.

**Figure 4:**
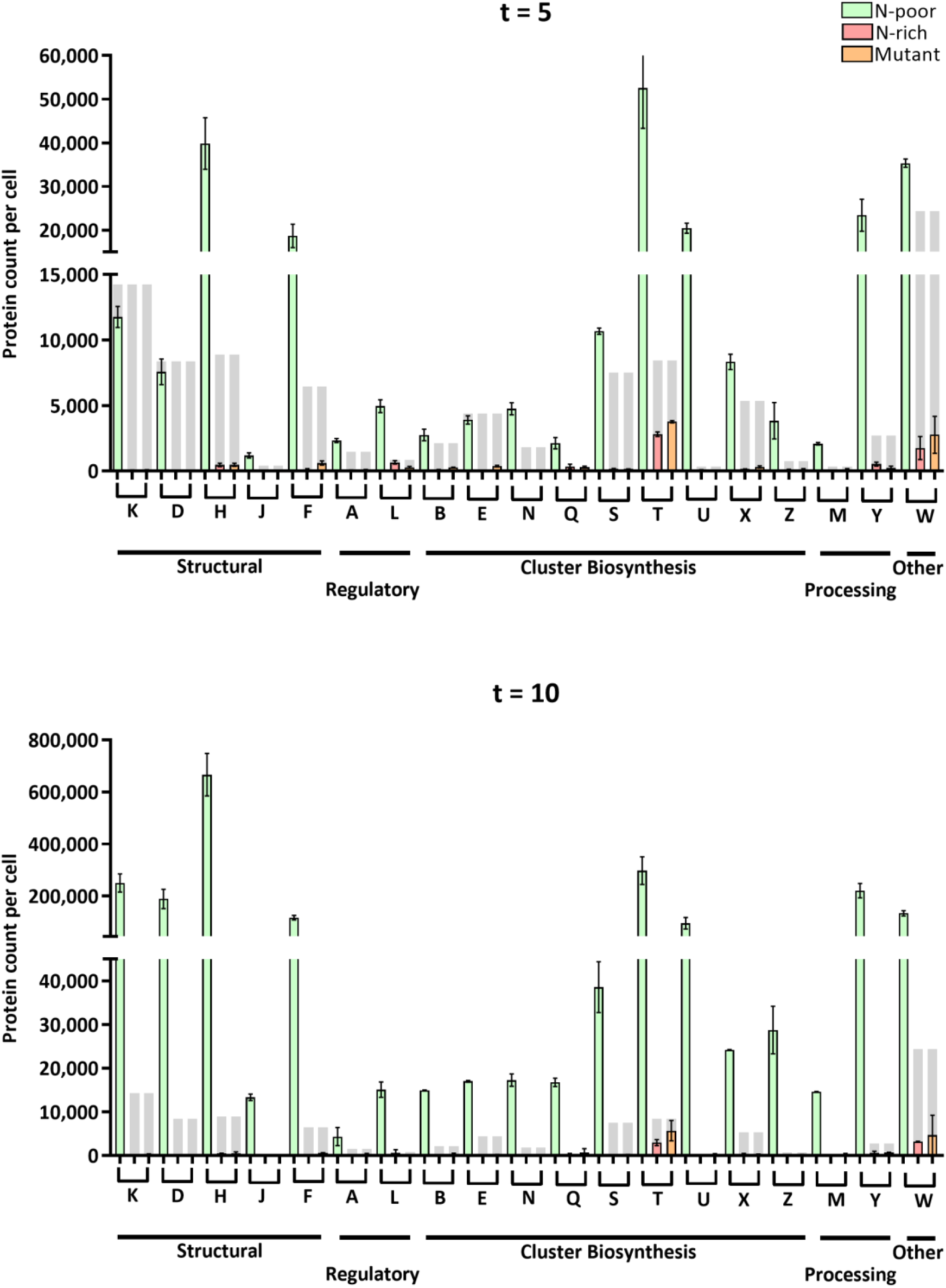
Nif protein stoichiometry at 5 and 10 hours after N sparging. Values per transition considered to be quantitatively robust were used to give an average per peptide, and these values were averaged in turn to give a value per Nif protein. Proteins are grouped by broad functionality, as indicated along the x axis. Y axes in both cases are split in order to better represent stoichiometry at lower protein counts. Error bars represent SEM over three biological replicates. The highest limit of quantification for each protein, based on figure 8a.II, is illustrated by a filled in light grey bar.

### QconCAT design and expression

Three QconCATs containing selected tryptic peptides from all Nif proteins. Nif proteins were grouped according to approximate intensity of corresponding MRM-MS spectra in samples. Since trypsin interacts with 3 to 4 residues around the scissile bond (15), each peptide was flanked with sequences of 3 amino acids on either side of the cleavage site, leading to 6 amino acid ‘linkers’, to ensure quantitative trypsin cleavage. ExPASy PeptideCutter (16) was used to confirm trypsin digestion efficiency at the cleavage sites within the native protein sequence and the QconCATs to be the same. After overexpression in *E. coli* BL21Δ*lys,arg*, each QconCAT was purified using Ni-affinity chromatography. Overexpression, molecular mass and urea solubilisation of Low, Medium and High QconCAT protein products were confirmed by SDS-PAGE (36.4, 37.6 and 29.7 kDa, respectively, (supplementary file). Table 2 shows the complete peptide list of the 3 QconCats.

**Table 2:**
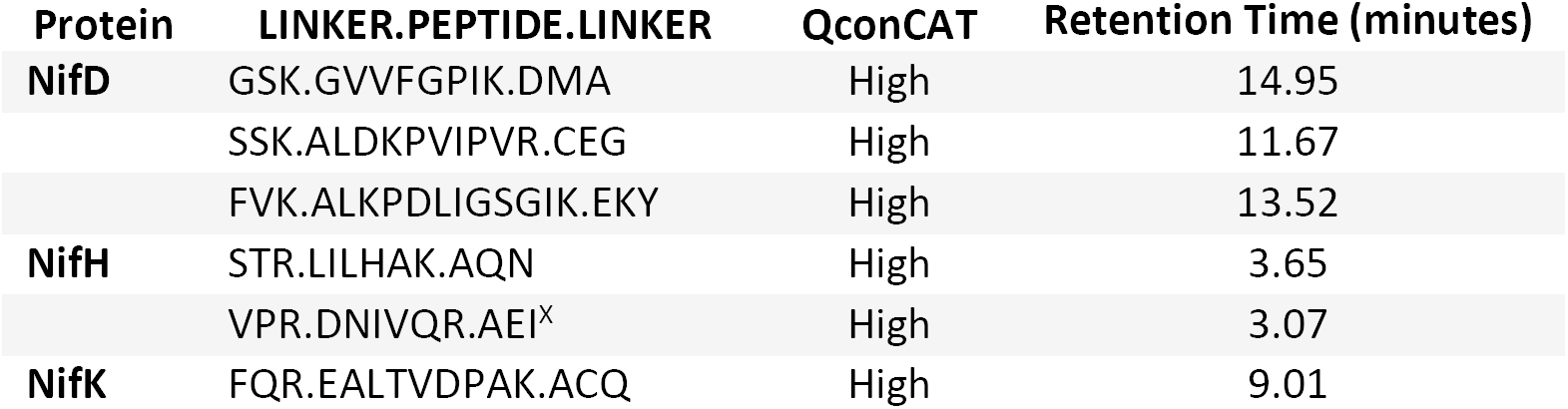
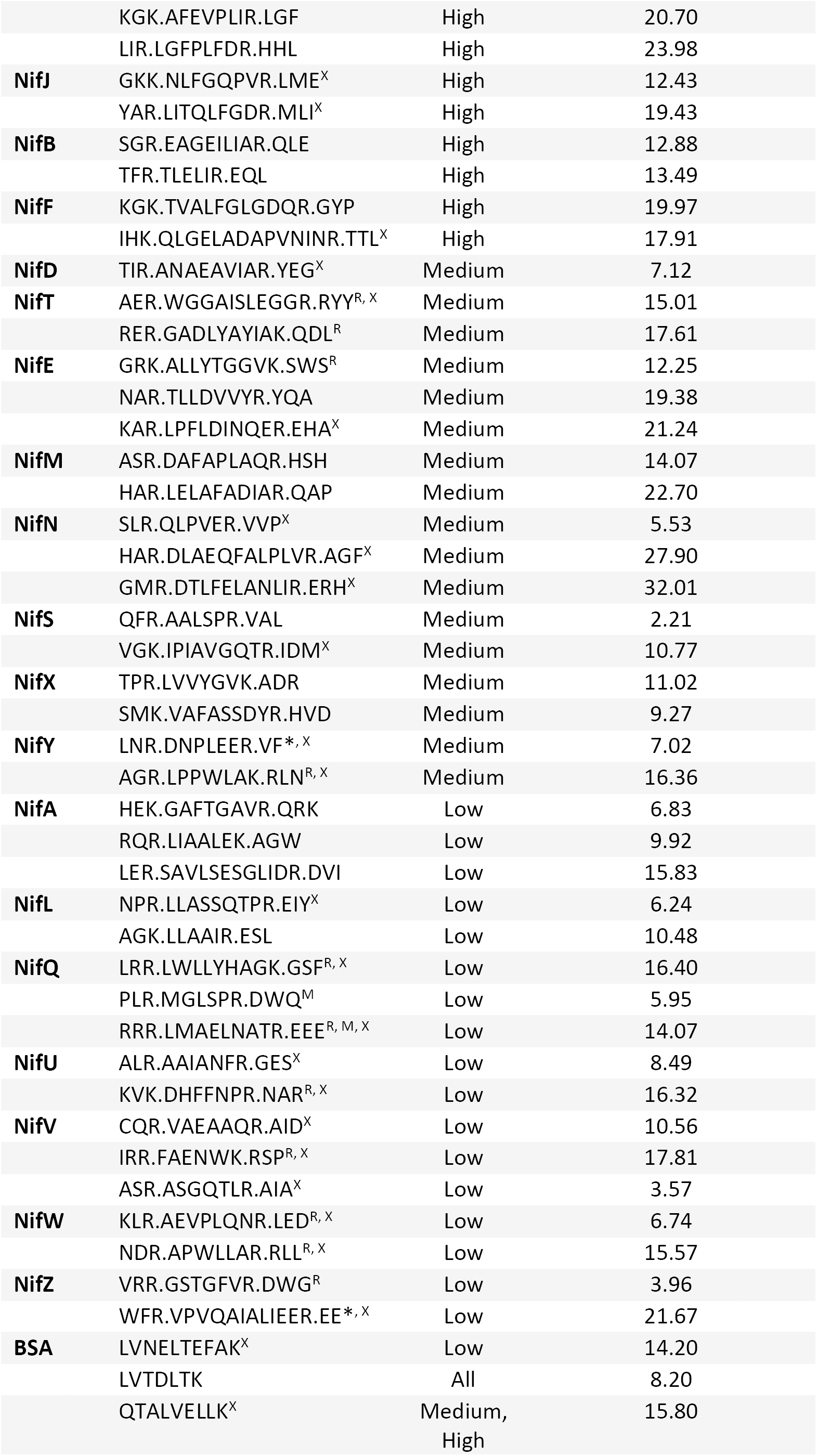
Final selection of Nif protein and BSA tryptic peptides, incorporated into three QconCATs (Low, Medium, High) based on approximate expression levels. Peptides are listed in the order in which they appear in the QconCAT. The linker amino acids on either side of the peptide represent the 5’ and 3’ natural flanking amino acids of each peptide within the Nif protein sequence. Peptides present at either end of the protein, and which are not bounded by three amino acids, have the missing amino acid denoted by a star. R, ragged end; M, methionine; X, contains residues susceptible to chemical or biological modifications. Retention times based on MRM-MS results are indicated and were considered during QconCAT design.

### Workflow validation

To establish the robustness of the QconCAT design expression pipeline prior to native Nif protein quantification, we interrogated general QconCAT LC and mass spectroscopic behaviour LC-MS (Appendix I), indicating that theoretical peptide selection yielded well resolved tryptic peptide spectra for each QconCAT (Figure 7a). We directly measured *in vivo* isotopic labelling efficiency following overexpression of QconCats in *E. coli* BL21Δ*lys,arg* in mutant strains, which showed 100% labelling efficiency, excluding the possibility of potential unspecific transamination in cells lacking both the specific transaminases for lysine and arginine.

Nif protein tryptic peptides were organised into Low, Medium and High QconCATs based on peak intensity (and thereby expression levels) during QconCAT design. The differences in the intensities were about ten-fold. 10-fold dilutions of each QconCAT were therefore prepared in order to determine biologically relevant concentrations, and samples prepared with identical native Nif protein concentration using a previously prepared N-fixing sample taken after 24 hours. The highest expressed Nif proteins were present at concentrations well above that of the undiluted High QconCAT (Figure c. I). Results for both the Medium and Low QconCATs (Figures c. II and III respectively), indicate that a 10-fold dilution of the QconCAT stock would lead to biologically relevant labelled Nif tryptic peptide concentrations.

### Validation of QconCAT-mediated quantification of Nif proteins by MRM-MS

The QconCAT method yields analytically rigorous data, and thereby accurate and precise quantitative measurement, by providing an isotopologue of the target analyte as an internal standard, to correlate signal intensity and analyte amount. Therefore, changes in analyte concentration should lead to proportional changes in signal, and thus a linear relationship between the two. Linearity can be compromised if the true protein concentration is not within the detection limit of the machinery used, or if the QconCAT is not an accurate measure of Nif protein concentration. Such a result could be observed if, for instance, if trypsin digestion efficiencies of QconCAT and native Nif protein were not comparable. We confirmed that all samples yielded > 90% trypsin digestion efficiency via SDS PAGE sypro-ruby staining and gel imaging, as we have done before (7). Both the native Nif proteins and the QconCAT were subject to linearity analysis, with signal calculated in terms of peak area. We confirmed that samples yielded > 90% trypsin Linearity of each QconCAT peptide standard was assayed within a complex bacterial matrix. The proportion of Nif protein (WT N-poor, 10 hours) in the matrix was varied from 100 to 0.001 % and QconCATs added at a constant concentration. Signal saturation was not apparent for any of the transitions as the slope of the dilution curves are consistently linear until 100 % Nif. All transitions for each peptide were assessed to determine the point below which the concentration to signal response ratio is no longer linear (‘quantification limit’). After assessing all transitions (Figure 2a. I.), those which showed no linearity anywhere over the whole % Nif range were removed (Figure 2a. II.) and were not included in further quantification analyses. Of the remaining transitions, linearity is broadly conserved in a 100-fold range from 1 to 100 % Nif protein, although the quantification limit of each transition, below which peak areas cannot be quantified with certainty, was noted for further analysis.

Conversely, QconCAT linearity within a constant bacterial protein matrix over a series of 2- and 5-fold dilutions was assayed. Most transitions for the High QconCAT were linear from a 0.1 to 2 dilution range (Figure 2b).

### Absolute quantification of Nif proteins in N-rich, N-poor and mutant K. oxytoca cultures

Quantification based on MRM-MS of Nif proteins was undertaken at two time points, early (t = 5) and late (t = 10), across the transition to diazotrophic growth, in a N-poor, N-rich and mutant culture (Figure 1). Purified QconCATs were added to complex protein samples derived from a time course experiments. Trypsin digestion efficiency determined by SDS-PAGE (supplementary file). The extracted ion chromatograms for unlabelled sample and labelled quantotypic peptide released from the QconCAT were used to calculate the amount of analyte, as protein count per cell. Quantitatively robust peptides were used in the quantification of Nif proteins. The peak area corresponding to the limit of quantification for each transition identified was calculated in terms of protein count per cell. Transitions giving protein counts below the limit of quantification were not included in further quantification. In general, the N-rich and mutant cultures were effective controls. The expression of some *nif* genes in the N-rich samples could be expected. The Δ*nifLA* strain should not express the NifL and NifA regulatory proteins, and hence should be *nif-*. However, some transitions corresponding to Nif protein tryptic peptides were visible in MRM-MS. Of these, however, the majority are below the limit set for quantitative accuracy, and the absence of other co-eluting transitions for the same peptide means their significance is questionable. Figure 3 shows the protein copy number per cell for the most highly abundant proteins (Nif H,D,K) and lowest abundance proteins (Nif J, F L, A). Abundances of all other twelve quantified proteins are shown in the supplementary file.

### Nif protein stoichiometries at early (t = 5) and late (t = 10) transition into N fixation of K. oxytoca

To determine the stoichiometries of Nif proteins absolute peptide quantities were averaged between based on counts of individual transitions. Stoichiometries provides information of the biochemical function of the nitrogenase, its assembly and regulation.

## Discussion

Since the publication of the QconCAT workflow (24), over 200 QconCATs have been used in a wide range of quantitative proteomic studies and analytical applications. Here we described the design, validation and quality control of a QconCAT approach with MRM-MS to measure 20 Nif proteins of *Klebsiella oxytoca*. Of the 20 Nif proteins included in the QconCATs, 19 passed the validation process and were quantified. The capability of MRM-MS to produce comprehensive datasets and quantify proteins over a wide dynamic range (32) allowed for quantification of proteins spanning more than 4 orders of magnitude; quantitative measures between 46 and 806630 proteins per cell were recorded.

No transitions corresponding to NifV peptides were visible in both analyte and QconCAT, leading to exclusion for NifV protein analysis. Peptide selection for NifV was problematic due to the very low intensity or non-symmetrical transitions and validation by EPI was not successful for NifV FAENWK and ASGQTLR. *In silico* digestion parameters had to be significantly broadened to select potential quantotypic peptides. For NifV FAENWK, and NifZ VPVQAIALIEER, peptide signal was observed for the QconCAT but not in bacterial samples. As such, quantification is in theory achievable by enrichment strategies.

### Reliability of quantification

Monitoring several peptides per protein greatly increases reliability of quantification, but can also highlight differences between peptides due to artefactual chemical modifications. These result in a fraction of the target peptide being converted to the modified form in an unpredictable manner (17). An Information Dependent Acquisition (IDA) MS method revealed prevalent modifications on NifD ALDKPVIPVR and NifK AFEVPLIR. Past quantification attempts using QconCATs also revealed discordant quantification data between a significant number of peptides, and suggested that two peptides per protein is inadequate for accurate quantification (15). Nonetheless, peptide modifications through e.g. post-translational modifications can be accurately measured, provided both a non-modified peptide and the modified peptide is included in the labelled standard as has been shown (18).

### Absolute Nif protein abundances and stoichiometries

The transition of *K. oxytoca* to diazotrophic growth requires a multiplicity of gene products, whose expression is highly coordinated. Steady state abundances of proteins is determined by function, and are based on, for instance, matching stoichiometry between proteins interacting within the same complexes (19). NifE and NifN contribute to FeMo-co biosynthesis by forming an α2β2 tetramer upon which FeMo-co is built, prior to insertion into the MoFe protein; hence, NifE and NifN are equimolar throughout the time course of this study (supplementary file). In addition, nitrogenase protein structure is such that the ratio expected between NifH, NifD and NifK is 2:1:1 (Figure 3). However, ratios decrease from approximately 5 to 3:1:1 between t = 5 and t = 10. As the cell transitions to diazotrophic growth, early NifH abundance is presumably required to also support both P-cluster and FeMo-co synthesis, perhaps explaining why it is present in an apparent excess with respect to known stoichiometry of the nitrogenase complex.

Within functional classes, changes in protein counts and stoichiometry were not always consistent, attesting to complex regulation of N fixation in *K. oxytoca*. There were, however, a few patterns that emerged. NifA and NifL counts were lower than most other Nif proteins, as is expected from regulatory proteins. NifL protein counts were consistently higher than NifA, perhaps because of the importance, due to high energy costs associated with N fixation, of a quick and effective sensing of unsuitable N fixation conditions. As expected, there is a high count of the structural NifH, D and K proteins, and 20 to 30-fold changes are observed between 5 and 10 hours, such that NifHDK accounts for 33 % of total protein at t = 10. As for the Nif proteins involved in cluster biosynthesis and processing, protein count increases from 4 to 10-fold between 5 and 10 hours, to support the increase in structural protein as *K. oxytoca* transitions into diazotrophic growth. The ratio between cluster biosynthesis/processing and structural Nif protein is higher at t = 5; early transition into diazotrophy presumably involves accumulation of biosynthetic proteins to a certain level, before upregulation of nitrogenase can be initiated.

Stoichiometry between biosynthetic proteins remained identical between 5 and 10 hours. Fold changes in biosynthetic/processing Nif protein count can be broadly split into a higher (NifQ, Z, B, M, Y and T) and lower (NifE, N, S, U, X) group. Protein counts for the lower group are at least half of their expected value at t = 10 if stoichiometry at t = 5 was to be preserved. Changes in stoichiometry could suggest that the final biosynthetic system allowing maximal N fixation has not yet been established.

Ratios between the nitrogenase components NifDK and Nif proteins involved in cluster biosynthesis and processing give insight into mature nitrogenase assembly mechanics. NifS, U, X, and W are all present at excess over the nascent nitrogenase protein at t = 5, but not at t = 10. These Nif proteins are presumably important in initial metabolic switching to diazotrophy. Indeed, previous studies have indicated that genes required for early nitrogenase biosynthesis (*nifU, nifS*) were expressed and turned off earlier than genes required for later stages of the pathway. Once fully functional N fixation was achieved, demand for biosynthetic components such as NifE, NifN and NifB decreased (8). It is probable that NifB and NifEN can perform multiple enzymatic turnovers *in vivo* to provide FeMo-co for an excess of NifDK.

Although NifT and NifY to NifDK ratio also decreases between t = 5 and t = 10, they are still required at substantial copy number at t = 10, with more than one NifT and NifY copy per NifDK. Whereas NifY has been characterised as a chaperone for the apo-MoFe protein, the exact role of NifT remains unknown. Given the 1:1:1 molar ratio between NifDK, NifT and NifY, it is possible that NifT also has a chaperone-like role in MoFe protein maturation. A summary scheme of Nif protein stoichiometries and their role in nitrogenase assembly and function is shown in Figure 5.

**Figure 5:**
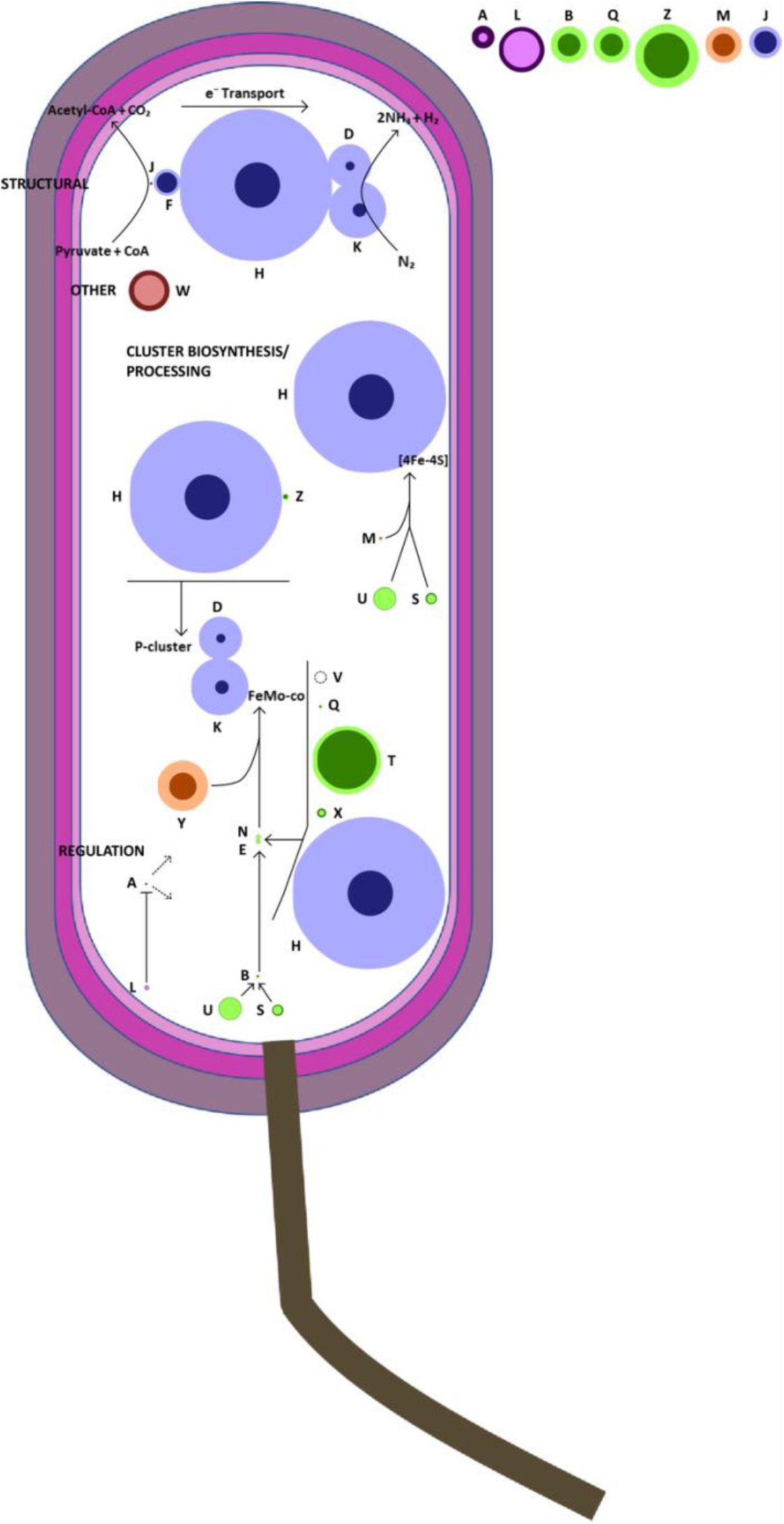
Representation of Nif protein stoichiometry in N-poor samples at t = 5 (dark circle) and t =10 (light circle), with illustration of cluster biosynthesis and nitrogen fixation pathways. Circle diameter is proportional to protein count per cell, with values at t = 10 halved in order to facilitate presentation. Nif proteins present at low counts are magnified 10-fold and displayed in top right corner of figure. Main classes of Nif proteins are indicated in capitals. Blue tones, structural; green tones, cluster biosynthesis; red tones, other; orange tones, processing; pink tones, regulation. Dotted lines indicate the regulation of *nif* gene expression by NifLA regulatory system. The role of NifV is indicated by a dotted circle, but no data was obtained for this protein.

Generally, we found that *nif* gene expression was very tightly regulated, with most Nif proteins below detection levels under N-rich conditions and up to hundreds of thousands of proteins per cell under diazotrophic growth. This stringent *nif* regulon control – apparently strictly NifLA dependent – differs from the nitrogen assimilation regulon controlled by NtrC in *E. coli*, where many genes are expressed despite ammonium starvation, implying complex and NtrC-independent regulatory mechanisms (36). Interestingly, both NifT and NifW are present in the N-rich and Δ*nifLA* control samples (Figure 10). It is possible an NifLA-independent regulatory mechanism of these two proteins exists, and their high values explained in part by involvement in other cellular processes. In studying the tolerance of nitrogenase activity to expression levels, NifT has been shown on multiple occasions to not have an effect on nitrogenase activity (20, 21), and is often absent from homologous clusters (38).

Previous studies on *Azotobacter vinelandii* have indicated that each *nif* operon yields similar mRNA levels for the genes it contains (8). At t = 10, a limited amount of agreement with this is observed. In *nifHDKTY* NifH is higher, in *nifENX* Nif X is higher, in *nifLA* NifL is higher than the remaining gene products of this operon, and in *nifUSVWZM* NifU and NifW are substantially higher than NifS, NifZ and NifM. However, a substantial number of regulatory processes occur after mRNA synthesis at the post-transcriptional, translational and protein degradation levels, and protein abundance reflects a dynamic balance between these processes. Mass spectrometric analysis, in particular used in conjunction with transcriptomics, can help establish to what extent each of these processes, as well as expression levels, contribute to regulation of cellular protein abundance (19). For instance, in *A. vinelandii*, cellular levels of NifB and NifEN proteins are controlled by the ClpX2 proteins by means of protein degradation, which is induced under diazotrophic conditions (22). Low NifEN and NifB may be the result of this regulatory system.

## Abbreviations

ATP: Adenosine Triphosphate
UAS: Upstream Activator Sequence
σ^54^: σ^54^ RNA polymerase
GS: Glutamine Synthetase
GOGAT: Glutamine Oxoglutarate Aminotransferase
GDH: Glutamate Dehydrogenase
UTase: Uridylyltransferase
UR: Uridylyl Removing
MS: Mass Spectrometry
ORF: Open Reading Frame
QconCAT: Artificial protein encoding a single ORF that is a concatenation of quantotypic tryptic peptides for a group of proteins under study
MRM-MS: Multiple Reaction Monitoring Mass Spectrometry
*m/z*: Mass/charge ratio
EPI: Enhanced Product Ion scan
BSA: Bovine Serum Albumin
LB: Luria-Bertani media
IPTG: Isopropyl β-D-1-thiogalactopyranoside
SDS-PAGE: Sodium Dodecyl Sulphate Polyacrylamide Gel Electrophoresis
TCEP: Tris(2-chloroethyl) Phosphate
UV: Ultraviolet
DC: Detergent Compatible
LC: Liquid Chromatography
IDA: Information Dependent Acquisition
N-poor: WT *K. oxytoca* strain M5a1 grown in 0.25 mM NH_4_
N-rich: WT *K. oxytoca* strain M5a1 grown in 10 mM NH_4_
Mutant: *K. oxytoca* M5a1 Δ*nifLA* grown in 0. 25 mM NH_4_
GC-MS: Gas Chromatography Mass Spectrometry
SEM: Standard Error of the Mean
WT: Wild-Type

