## Supplementary File for "A multiplexed work-flow for absolute quantification of *Klebsiella oxytoca* Nif proteins"

### Absolute quantification of *Klebsiella oxytoca* Nif proteins using a synthetic QconCAT strategy

Supplementary Table 1: *Klebsiella oxytoca nif* genes and functions.

| Gene | Identity and/or role of gene product |
|---|---|
| **Structural** | |
| ***nifH*** | Fe protein, a homodimer of nitrogenase. NifH is an obligate electron donor to the MoFe protein and is required for FeMo-co biosynthesis as well as apo-MoFe protein maturation. |
| ***nifD*** | α chain of the MoFe protein, an α2β2 tetramer of nitrogenase. Substrate reduction at FeMo-co occurs within the α subunit. |
| ***nifK*** | β chain of the MoFe protein. P clusters are present at the αβ interfaces. |
| ***nifJ*** | Pyruvate:flavodoxin (ferrodoxin) oxidoreductase. Electron donor to the Fe protein, provided by oxidative decarboxylation of pyruvate to acetyl-CoA. |
| ***nifF*** | Flaxodoxin. Electron donor to the Fe protein. |
| **Regulatory** | |
| ***nifA*** | $\sigma^{54}$-dependent activator of *nif* gene transcription. Co-transcribed with *nifL*. |
| ***nifL*** | *nif* gene negative regulator. Senses redox and fixed nitrogen status to regulate nitrogen fixation by modulating activity of NifA. |
| **Cluster Biosynthesis** | |
| ***nifQ*** | Involved in FeMo-co biosynthesis. Proposed to function in early $MoO_4^{-2}$ processing. |
| ***nifB*** | Involved in FeMo-co biosynthesis. NifB-co is a specific Fe and S donor to FeMo-co, and is transferred to the NifEN complex where it is processed to FeMo-co. |
| ***nifN*** | Involved in FeMo-co biosynthesis. Forms an α2β2 tetramer with NifE which has been proposed to act as a scaffold upon which FeMo-co is built and then inserted into the MoFe protein. |
| ***nifE*** | As above. |
| ***nifV*** | Homocitrate synthase. Involved in FeMo-co biosynthesis although exact role remains unknown. |
| ***nifS*** | Cysteine desulphurase. Involved in FeS (both [8Fe-7S] and [4Fe-4S]) cluster biosynthesis and repair by mobilization of S. |
| ***nifZ*** | Involved in MoFe protein maturation. Suggested to be specifically required for the formation of the second P-cluster. |
| ***nifU*** | Involved in FeS cluster biosynthesis. Homodimer of two subunits each containing a 2Fe-2S cluster, proposed to have a redox function involving release of Fe for FeS cluster biosynthesis. |
| ***nifX*** | Involved in FeMo-co biosynthesis. Appears to be associated with mature FeMo-co and accumulates an FeSMo-containing precursor. |
| ***nifT*** | Involved in FeMo-co biosynthesis. Exact role remains unknown. |
| **Processing** | |
| ***nifM*** | Accessory protein for NifH, required for NifH maturation. |
| ***nifY*** | Chaperone for the apo-MoFe protein. |
| **Other** | |
| ***nifW*** | Proposed to be involved in maturation or stability of the MoFe protein. Other studies have proposed a role in protection of MoFe protein from oxygen damage. |

Supplementary Figure 1: Regulation of nitrogen assimilation and nitrogen fixation of Klebsiella oxytoca

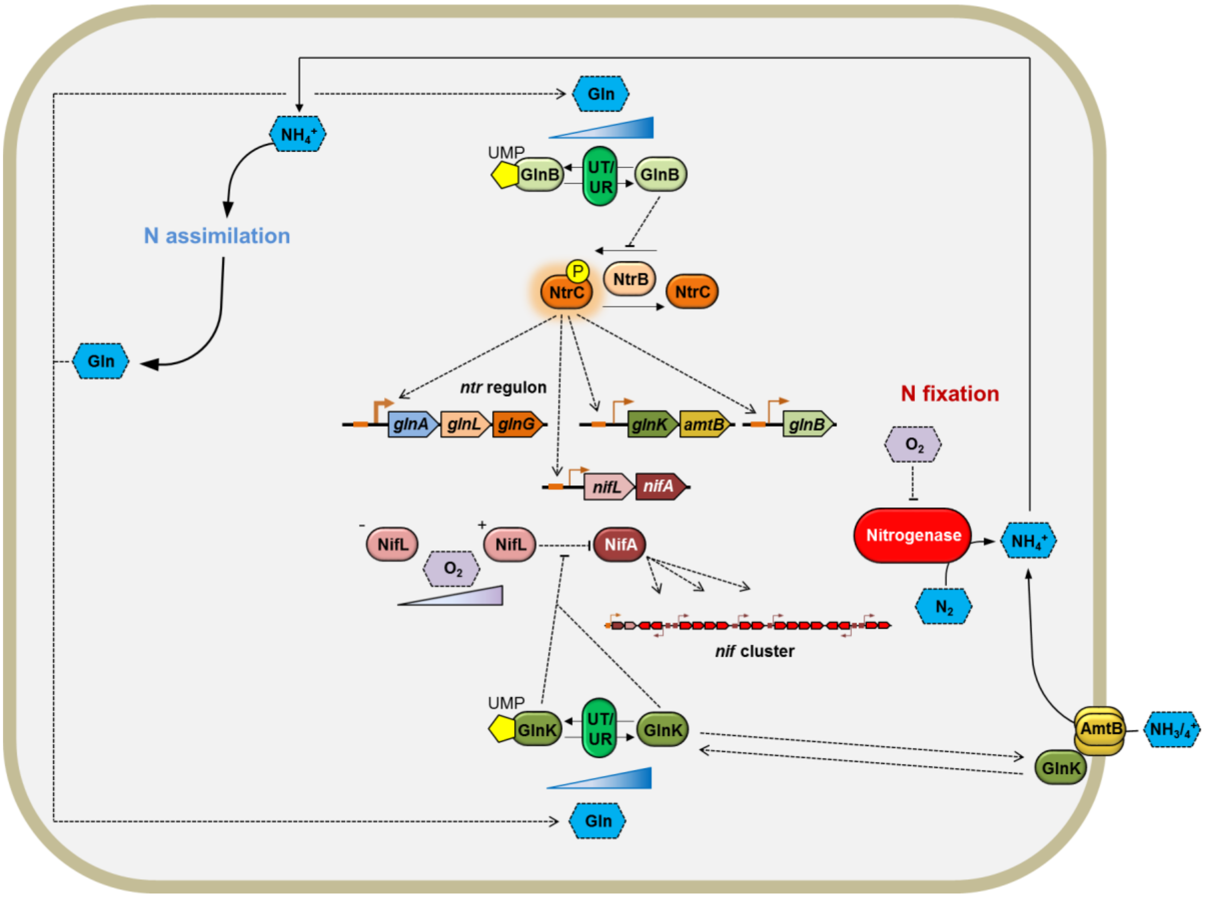


**Figure 3: Regulation of nitrogen fixation in *K. oxytoca.*** Assembly and synthesis of the nitrogenase complex and subsequent dinitrogen reduction is stringently controlled and regulated on multiple levels, and in response to multiple environmental cues. Nitrogen assimilation acts as an input into the control of nitrogen fixation, primarily by glutamine sensing. Genes and corresponding gene products are of the same colour. Genes are illustrated by large arrows, gene products by ovals, metabolites/chemical species by hexagons. PII, GlnB. Diagram made by Chris Waite (Imperial College London).

Supplementary Figure 2

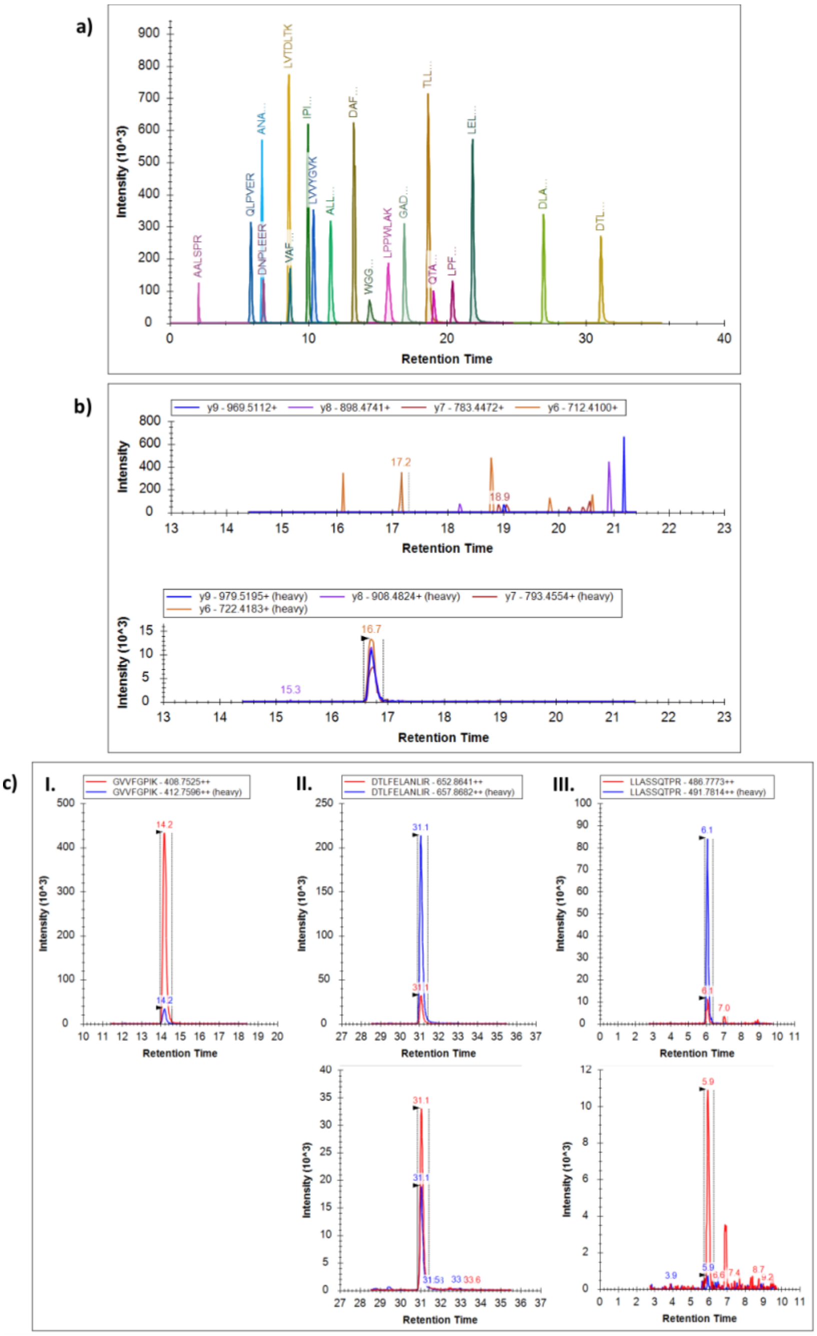


**Supplementary Figure 2: MRM-MS profiles for Nif protein QconCATs.** (a) Spectrum generated by MRM-MS corresponding to tryptic peptides in the Medium QconCAT, with peaks labelled for each tryptic peptide. Peptides longer than 7 amino acids are indicated by their first three letters only. (b) Spectrum of QLGELADAPVNINR. Top, spectrum generated from light transitions, a peak in which would correspond to unlabelled QconCAT. Bottom, spectrum generated from heavy transitions, corresponding to labelled QconCAT. Names and m/z for each transition are indicated in each legend. (c) QconCATs (blue, heavy) were diluted 0 (High), 10 (Medium) and 100 (Low) times with the same concentration of sample (red, light) (as indicated in the legends). Results show the spectrum of a selected peptide from each QconCAT, with heavy and light signal overlain, the sample with undiluted QconCAT at the top and the diluted QconCAT at the bottom. A 10-fold dilution of the Low QconCAT should lead to peak intensity of approximately 8, which is much more biologically relevant. Images exported from Skyline.

Supplementary Figure 3: Absolute quantification of Nif proteins and limits of quantification over time and under different conditions.

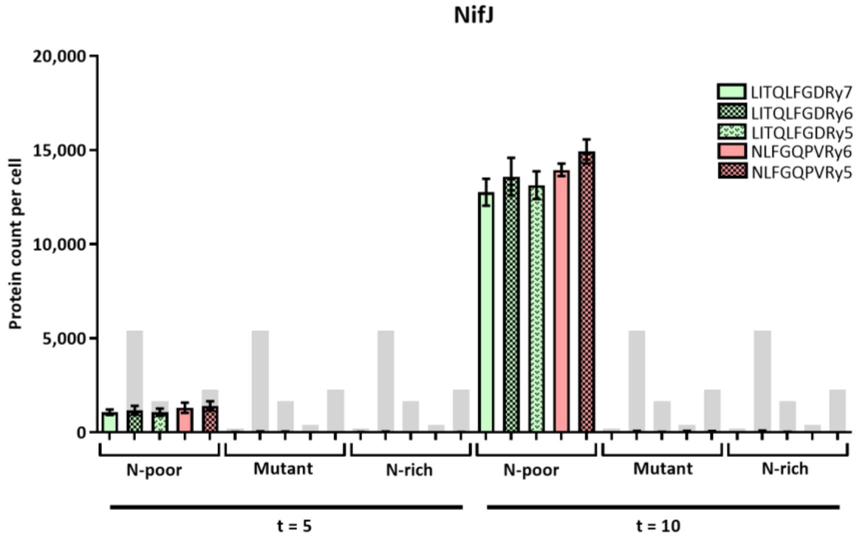
NifJ
Protein count per cell
20,000
15,000
10,000
5,000
0
LITQLFGDRy7
LITQLFGDRy6
LITQLFGDRy5
NLFGQPVRy6
NLFGQPVRy5
N-poor
Mutant
N-rich
N-poor
Mutant
N-rich
t = 5
t = 10

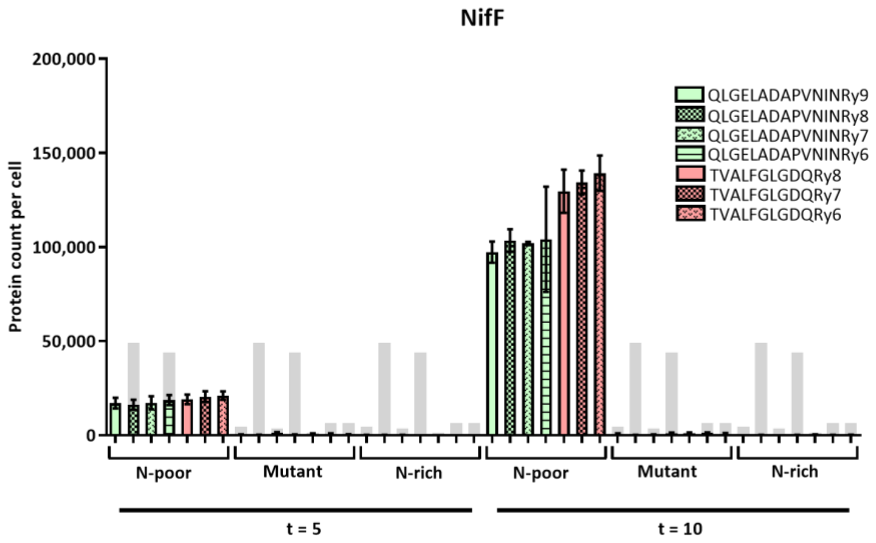
NifF
Protein count per cell
200,000
150,000
100,000
50,000
0
QLGELADAPVNINRy9
QLGELADAPVNINRy8
QLGELADAPVNINRy7
QLGELADAPVNINRy6
TVALFGLGDQRy8
TVALFGLGDQRy7
TVALFGLGDQRy6
N-poor
Mutant
N-rich
N-poor
Mutant
N-rich
t = 5
t = 10

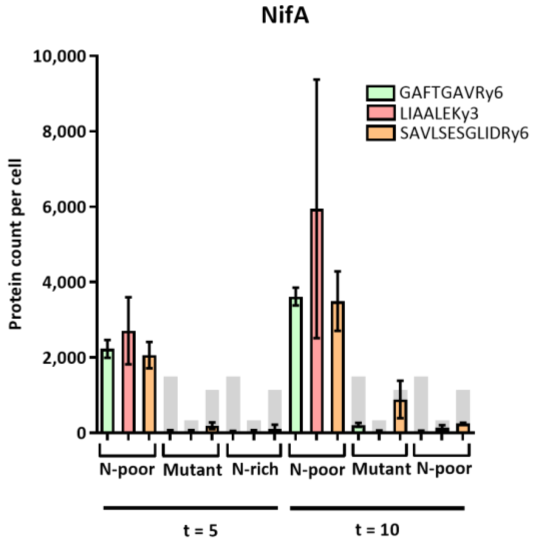
NifA
Protein count per cell
10,000
8,000
6,000
4,000
2,000
0
GAFTGAVRy6
LIAALEKy3
SAVLSESGLIDRy6
N-poor
Mutant
N-rich
N-poor
Mutant
N-poor
t = 5
t = 10

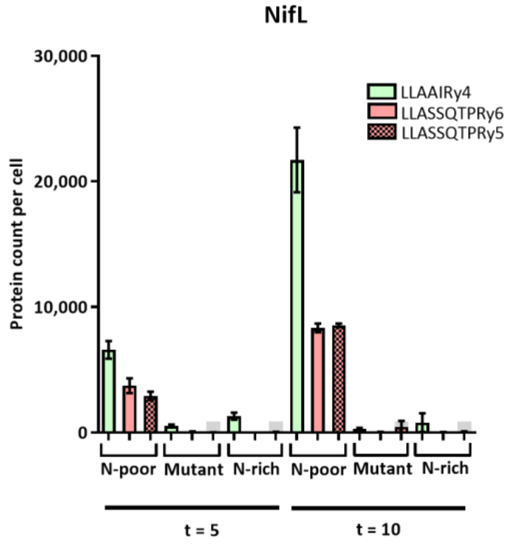
NifL
Protein count per cell
30,000
20,000
10,000
0
LLAAIRy4
LLASSQTPRy6
LLASSQTPRy5
N-poor
Mutant
N-rich
N-poor
Mutant
N-rich
t = 5
t = 10

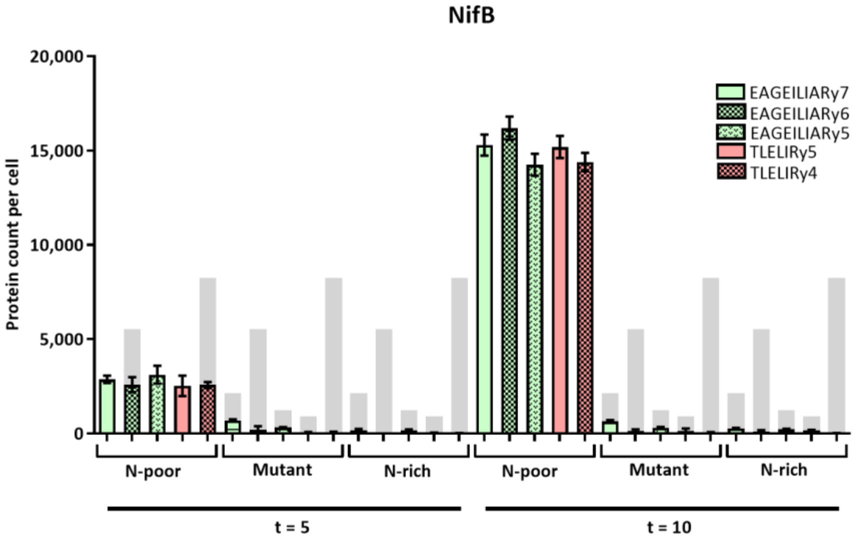
NifB
Protein count per cell
20,000
15,000
10,000
5,000
0
EAGEILIARy7
EAGEILIARy6
EAGEILIARy5
TLELIRy5
TLELIRy4
N-poor
Mutant
N-rich
N-poor
Mutant
N-rich
t = 5
t = 10

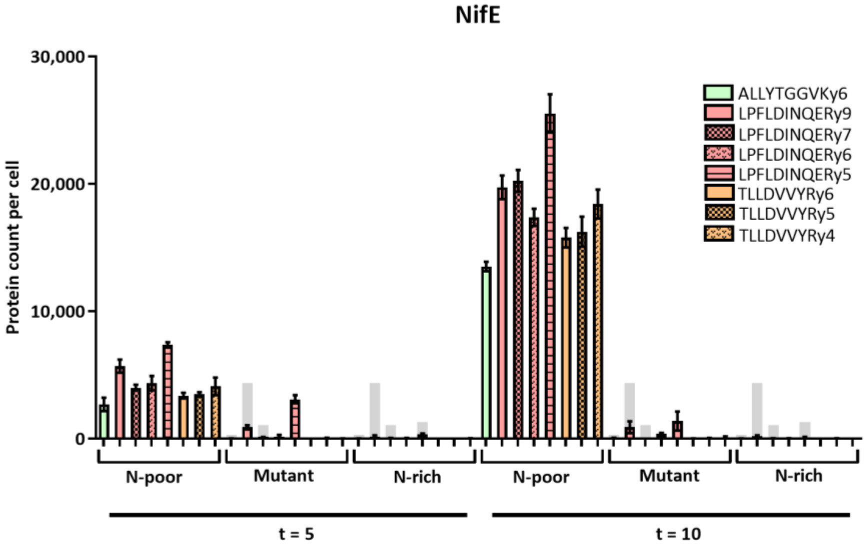
NifE
Protein count per cell
30,000
20,000
10,000
0
ALLYTGGVKy6
LPFLDINQERy9
LPFLDINQERy7
LPFLDINQERy6
LPFLDINQERy5
TLLDVVYRy6
TLLDVVYRy5
TLLDVVYRy4
N-poor
Mutant
N-rich
N-poor
Mutant
N-rich
t = 5
t = 10

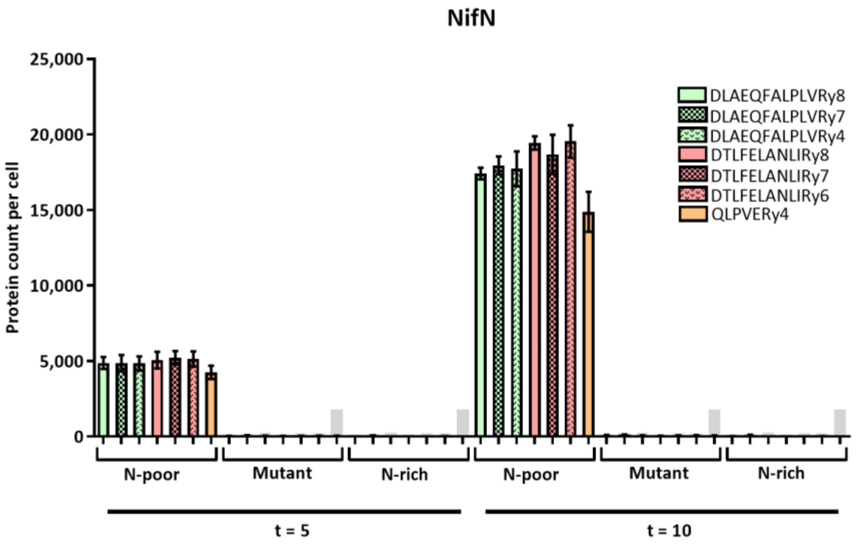
NifN
Protein count per cell
25,000
20,000
15,000
10,000
5,000
0
DLAEQFALPLVRy8
DLAEQFALPLVRy7
DLAEQFALPLVRy4
DTLFELANLIRy8
DTLFELANLIRy7
DTLFELANLIRy6
QLPVERy4
N-poor
Mutant
N-rich
N-poor
Mutant
N-rich
t = 5
t = 10

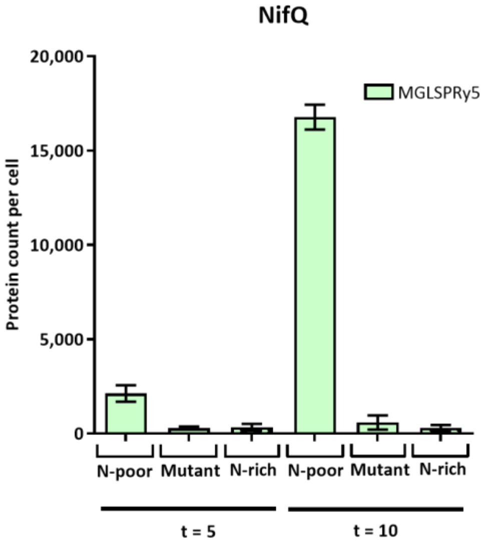
NifQ
MGLSPRy5
Protein count per cell
20,000
15,000
10,000
5,000
0
N-poor
Mutant
N-rich
N-poor
Mutant
N-rich
t = 5
t = 10

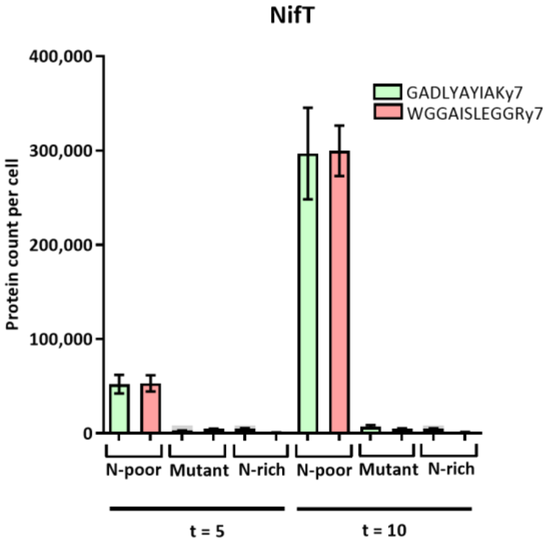
NifT
GADLYAYIAKy7
WGGAISLEGGRy7
Protein count per cell
400,000
300,000
200,000
100,000
0
N-poor
Mutant
N-rich
N-poor
Mutant
N-rich
t = 5
t = 10

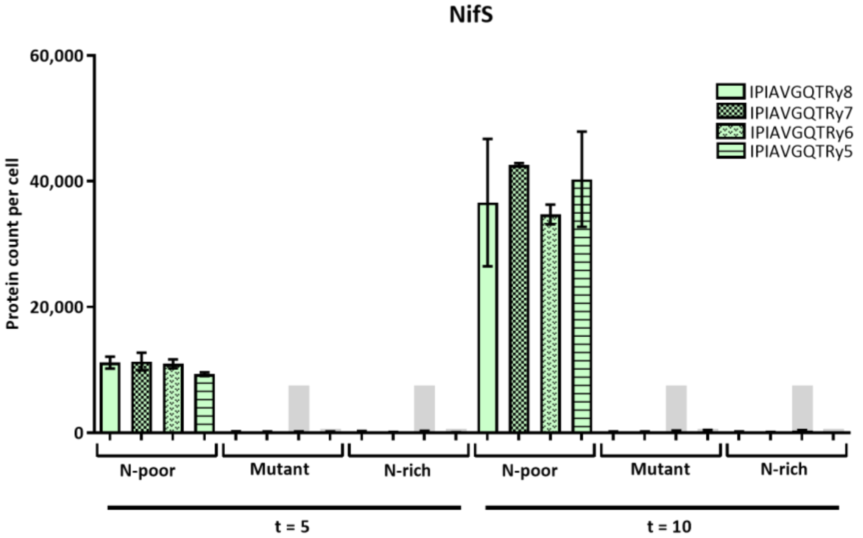
NifS
IPIAVGQTRy8
IPIAVGQTRy7
IPIAVGQTRy6
IPIAVGQTRy5
Protein count per cell
60,000
40,000
20,000
0
N-poor
Mutant
N-rich
N-poor
Mutant
N-rich
t = 5
t = 10

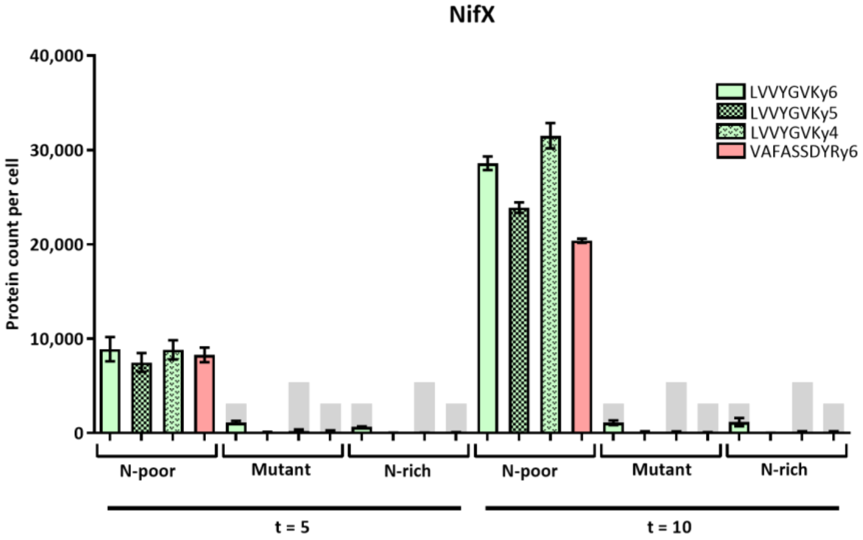
NifX
LVVYGVKy6
LVVYGVKy5
LVVYGVKy4
VAFASSDYRy6
Protein count per cell
40,000
30,000
20,000
10,000
0
N-poor
Mutant
N-rich
N-poor
Mutant
N-rich
t = 5
t = 10

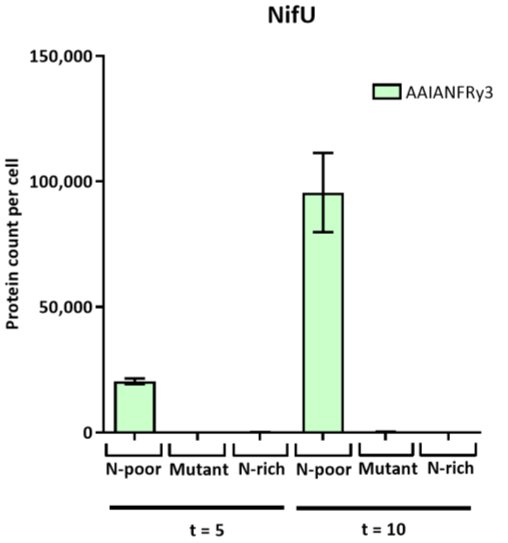
NifU
150,000
100,000
50,000
0
Protein count per cell
AAIANFRy3
N-poor
Mutant
N-rich
N-poor
Mutant
N-rich
t = 5
t = 10

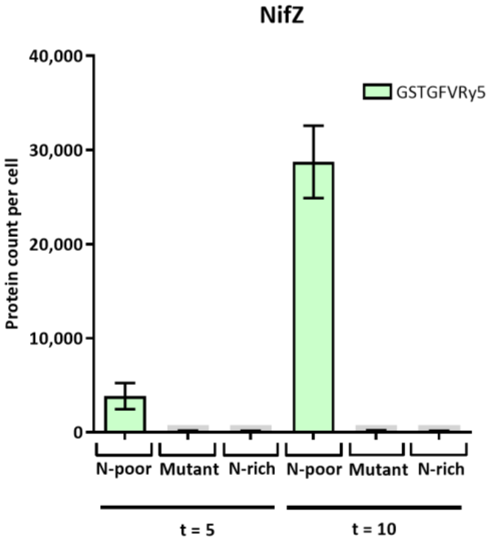
NifZ
40,000
30,000
20,000
10,000
0
Protein count per cell
GSTGFVRy5
N-poor
Mutant
N-rich
N-poor
Mutant
N-rich
t = 5
t = 10

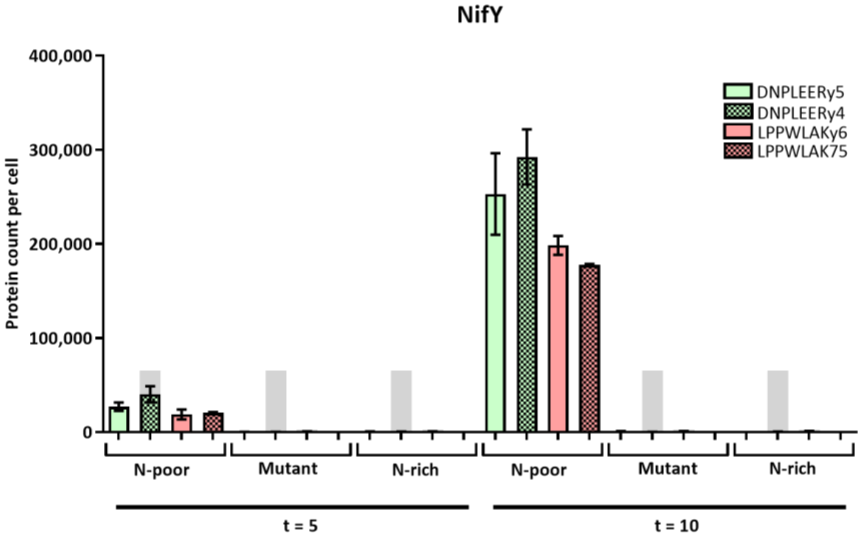
NifY
400,000
300,000
200,000
100,000
0
Protein count per cell
DNPLEERy5
DNPLEERy4
LPPWLAKy6
LPPWLAK75
N-poor
Mutant
N-rich
N-poor
Mutant
N-rich
t = 5
t = 10

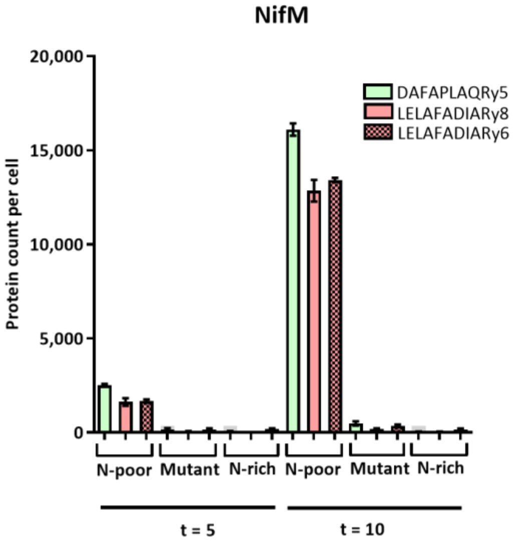
NifM
20,000
15,000
10,000
5,000
0
Protein count per cell
DAFAPLAQRy5
LELAFADIARy8
LELAFADIARy6
N-poor
Mutant
N-rich
N-poor
Mutant
N-rich
t = 5
t = 10

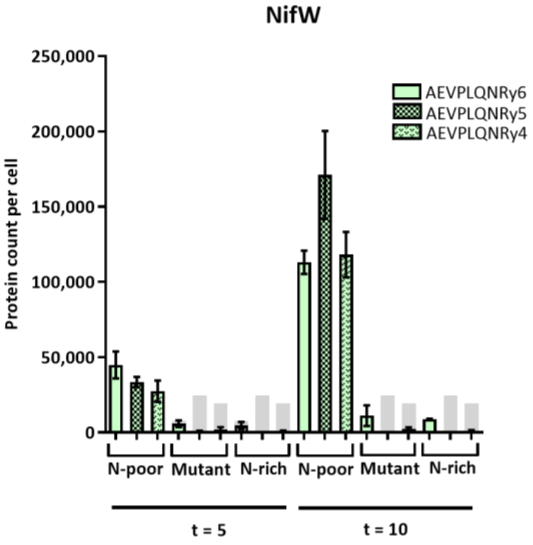
NifW
250,000
200,000
150,000
100,000
50,000
0
Protein count per cell
AEVPLQNRy6
AEVPLQNRy5
AEVPLQNRy4
N-poor
Mutant
N-rich
N-poor
Mutant
N-rich
t = 5
t = 10

**Supplementary Figure 3 (previous 5 pages): Protein count per cell for all individual Nif proteins in K. *oxytoca* M5a1 grown in 0.25 (N-poor) and 10 (N-rich) mM $NH_4$, and *K. oxytoca* M5a1 Δ*nifLA* (mutant) grown in 0.25 mM $NH_4$, at both 5 and 10 hours after N sparging.** All transitions for each peptide are shown, and limit of quantification illustrated by a filled in light grey bar. Error bars represent SEM over three biological replicates. Note that y-axis scale varies.

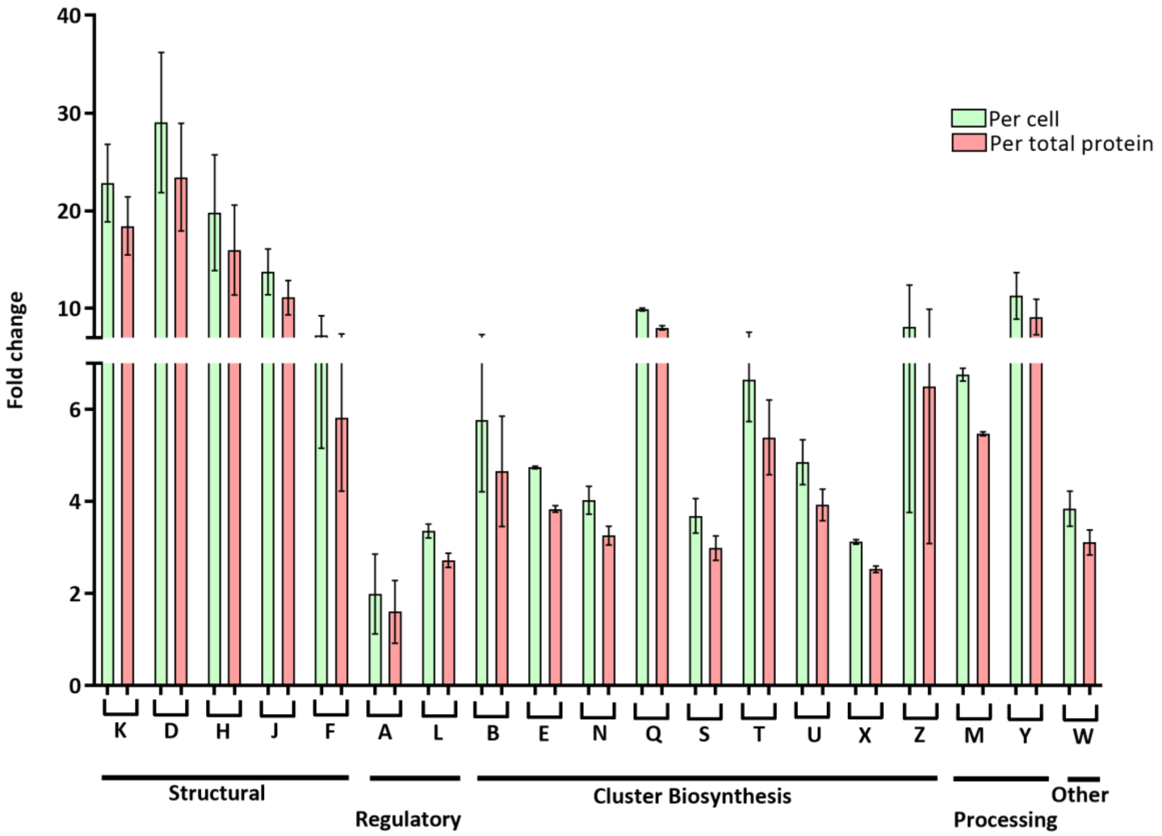


**Supplementary Figure 4: Fold changes of Nif protein counts between t =5 and t =10, calculated per cell and per total protein.** General function of each Nif protein is indicated along the x-axis. The y-axis is split in order to provide fold-change information at lower values. Error bars indicate SEM over 2 biological replicates.

Supplementary Figure 5:

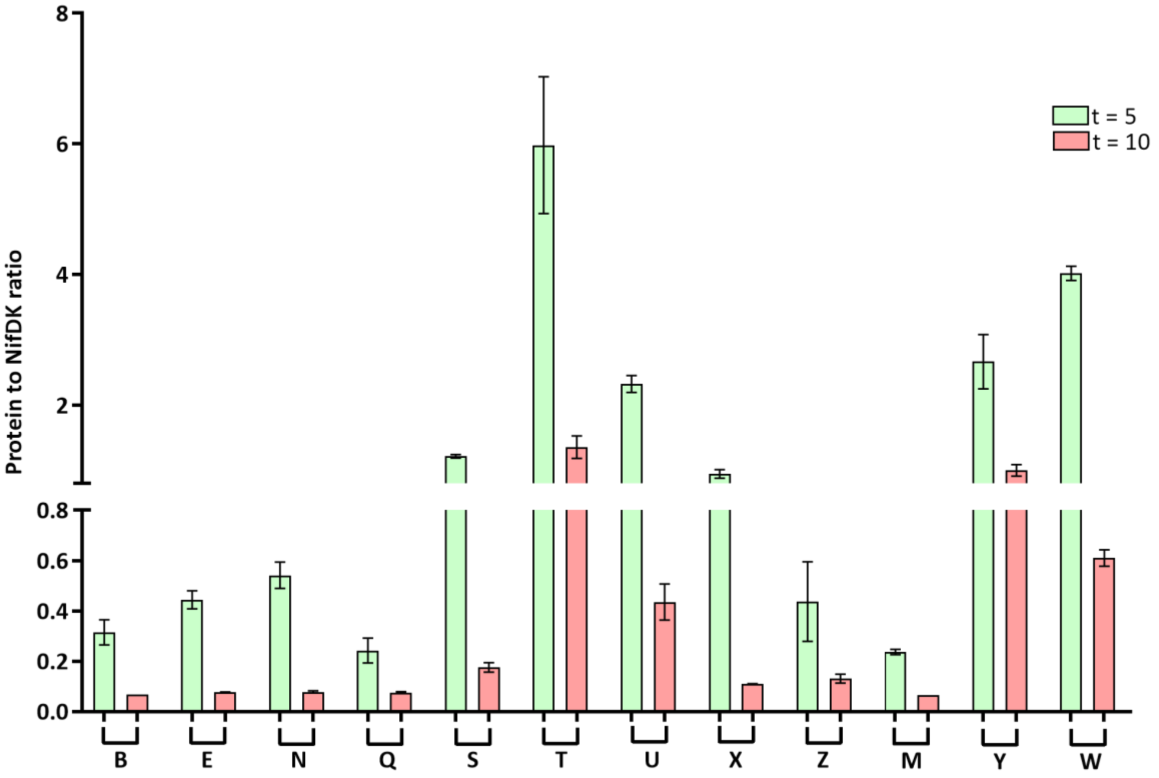


**Supplementary Figure 5: Ratio of each Nif protein involved in nitrogenase cluster biosynthesis and processing to NifDK, at t = 5 and t = 10.** Nif protein is indicated along the x-axis. Error bars represent SEM over three biological replicates. Note that y-axis is split at 0.8.

Supplementary Table 2: List of all transitions and selected transitions used in final analysis

| Q1 mass | Q3 mass | Retention time | Transition identifier | Group name | W | DP | CE |
|---|---|---|---|---|---|---|---|
| 586.3261 | 603.3228 | 14.2 | e3.BSA.LVNELTEFAK.+2y5.heavy | BSA-2 | 0.5 | 73.6 | 29.8 |
| 582.319 | 595.3086 | 14.2 | e3.BSA.LVNELTEFAK.+2y5.light | BSA-2 | 0.5 | 73.6 | 29.8 |
| 586.3261 | 716.4069 | 14.2 | e3.BSA.LVNELTEFAK.+2y6.heavy | BSA-2 | 0.5 | 73.6 | 29.8 |
| 582.319 | 708.3927 | 14.2 | e3.BSA.LVNELTEFAK.+2y6.light | BSA-2 | 0.5 | 73.6 | 29.8 |
| 586.3261 | 845.4495 | 14.2 | e3.BSA.LVNELTEFAK.+2y7.heavy | BSA-2 | 0.5 | 73.6 | 29.8 |
| 582.319 | 837.4353 | 14.2 | e3.BSA.LVNELTEFAK.+2y7.light | BSA-2 | 0.5 | 73.6 | 29.8 |
| 394.7181 | 412.2542 | 6.3 | e3.NifA.GAFTGAVR.+2y4.heavy | NifA-1 | 4 | 59.5 | 22.9 |
| 389.7139 | 402.2459 | 6.3 | e3.NifA.GAFTGAVR.+2y4.light | NifA-1 | 4 | 59.5 | 22.9 |
| 394.7181 | 513.3019 | 6.3 | e3.NifA.GAFTGAVR.+2y5.heavy | NifA-1 | 4 | 59.5 | 22.9 |
| 389.7139 | 503.2936 | 6.3 | e3.NifA.GAFTGAVR.+2y5.light | NifA-1 | 4 | 59.5 | 22.9 |
| **394.7181** | **660.3703** | **6.3** | **e3.NifA.GAFTGAVR.+2y6.heavy** | **NifA-1** | **4** | **59.5** | **22.9** |
| **389.7139** | **650.362** | **6.3** | **e3.NifA.GAFTGAVR.+2y6.light** | **NifA-1** | **4** | **59.5** | **22.9** |
| **383.2516** | **397.2537** | **9.7** | **e3.NifA.LIAALEK.+2y3.heavy** | **NifA-2** | **2** | **58.8** | **22.5** |
| **379.2445** | **389.2395** | **9.7** | **e3.NifA.LIAALEK.+2y3.light** | **NifA-2** | **2** | **58.8** | **22.5** |
| 383.2516 | 468.2908 | 9.7 | e3.NifA.LIAALEK.+2y4.heavy | NifA-2 | 2 | 58.8 | 22.5 |
| 379.2445 | 460.2766 | 9.7 | e3.NifA.LIAALEK.+2y4.light | NifA-2 | 2 | 58.8 | 22.5 |
| 383.2516 | 539.3279 | 9.7 | e3.NifA.LIAALEK.+2y5.heavy | NifA-2 | 2 | 58.8 | 22.5 |
| 379.2445 | 531.3137 | 9.7 | e3.NifA.LIAALEK.+2y5.light | NifA-2 | 2 | 58.8 | 22.5 |
| **628.8397** | **670.3758** | **14.5** | **e3.NifA.SAVLSESGLIDR.+2y6.heavy** | **NifA-3** | **4** | **76.6** | **31.3** |
| **623.8355** | **660.3675** | **14.5** | **e3.NifA.SAVLSESGLIDR.+2y6.light** | **NifA-3** | **4** | **76.6** | **31.3** |
| 628.8397 | 799.4184 | 14.5 | e3.NifA.SAVLSESGLIDR.+2y7.heavy | NifA-3 | 4 | 76.6 | 31.3 |
| 623.8355 | 789.4101 | 14.5 | e3.NifA.SAVLSESGLIDR.+2y7.light | NifA-3 | 4 | 76.6 | 31.3 |
| 628.8397 | 886.4504 | 14.5 | e3.NifA.SAVLSESGLIDR.+2y8.heavy | NifA-3 | 4 | 76.6 | 31.3 |
| 623.8355 | 876.4421 | 14.5 | e3.NifA.SAVLSESGLIDR.+2y8.light | NifA-3 | 4 | 76.6 | 31.3 |
| 333.7305 | 369.2484 | 9.4 | e3.NifL.LLAAIR.+2y3.heavy | NifL-2 | 1 | 55.1 | 20.7 |
| 328.7263 | 359.2401 | 9.4 | e3.NifL.LLAAIR.+2y3.light | NifL-2 | 1 | 55.1 | 20.7 |
| **333.7305** | **440.2855** | **9.4** | **e3.NifL.LLAAIR.+2y4.heavy** | **NifL-2** | **1** | **55.1** | **20.7** |
| **328.7263** | **430.2772** | **9.4** | **e3.NifL.LLAAIR.+2y4.light** | **NifL-2** | **1** | **55.1** | **20.7** |
| 333.7305 | 553.3696 | 9.4 | e3.NifL.LLAAIR.+2y5.heavy | NifL-2 | 1 | 55.1 | 20.7 |
| 328.7263 | 543.3613 | 9.4 | e3.NifL.LLAAIR.+2y5.light | NifL-2 | 1 | 55.1 | 20.7 |
| 491.7814 | 511.2862 | 5.9 | e3.NifL.LLASSQTPR.+2y4.heavy | NifL-1 | 4 | 66.6 | 26.4 |
| 486.7773 | 501.278 | 5.9 | e3.NifL.LLASSQTPR.+2y4.light | NifL-1 | 4 | 66.6 | 26.4 |
| **491.7814** | **598.3183** | **5.9** | **e3.NifL.LLASSQTPR.+2y5.heavy** | **NifL-1** | **4** | **66.6** | **26.4** |
| **486.7773** | **588.31** | **5.9** | **e3.NifL.LLASSQTPR.+2y5.light** | **NifL-1** | **4** | **66.6** | **26.4** |
| **491.7814** | **685.3503** | **5.9** | **e3.NifL.LLASSQTPR.+2y6.heavy** | **NifL-1** | **4** | **66.6** | **26.4** |
| **486.7773** | **675.342** | **5.9** | **e3.NifL.LLASSQTPR.+2y6.light** | **NifL-1** | **4** | **66.6** | **26.4** |
| 514.7753 | 584.339 | 10 | e3.NifQ.LMAELNATR.+2y5.heavy | NifQ-3 | 4 | 68.3 | 27.2 |
| 509.7711 | 574.3307 | 10 | e3.NifQ.LMAELNATR.+2y5.light | NifQ-3 | 4 | 68.3 | 27.2 |
| 514.7753 | 713.3816 | 10 | e3.NifQ.LMAELNATR.+2y6.heavy | NifQ-3 | 4 | 68.3 | 27.2 |
| 509.7711 | 703.3733 | 10 | e3.NifQ.LMAELNATR.+2y6.light | NifQ-3 | 4 | 68.3 | 27.2 |
| 514.7753 | 784.4187 | 10 | e3.NifQ.LMAELNATR.+2y7.heavy | NifQ-3 | 4 | 68.3 | 27.2 |
| 509.7711 | 774.4104 | 10 | e3.NifQ.LMAELNATR.+2y7.light | NifQ-3 | 4 | 68.3 | 27.2 |
| 554.8233 | 583.3078 | 15.7 | e3.NifQ.LWLLYHAGK.+2y5.heavy | NifQ-1 | 4 | 71.3 | 28.7 |

| | | | | | | | |
|---|---|---|---|---|---|---|---|
| 550.8162 | 575.2936 | 15.7 | e3.NifQ.LWLLYHAGK.+2y5.light | NifQ-1 | 4 | 71.3 | 28.7 |
| 554.8233 | 696.3919 | 15.7 | e3.NifQ.LWLLYHAGK.+2y6.heavy | NifQ-1 | 4 | 71.3 | 28.7 |
| 550.8162 | 688.3777 | 15.7 | e3.NifQ.LWLLYHAGK.+2y6.light | NifQ-1 | 4 | 71.3 | 28.7 |
| 554.8233 | 809.4759 | 15.7 | e3.NifQ.LWLLYHAGK.+2y7.heavy | NifQ-1 | 4 | 71.3 | 28.7 |
| 550.8162 | 801.4618 | 15.7 | e3.NifQ.LWLLYHAGK.+2y7.light | NifQ-1 | 4 | 71.3 | 28.7 |
| 370.218 | 420.2445 | 15.7 | e3.NifQ.LWLLYHAGK.+3y4.heavy | NifQ-1 | 4 | 57.9 | 17.6 |
| 367.5466 | 412.2303 | 15.7 | e3.NifQ.LWLLYHAGK.+3y4.light | NifQ-1 | 4 | 57.9 | 17.6 |
| 370.218 | 583.3078 | 15.7 | e3.NifQ.LWLLYHAGK.+3y5.heavy | NifQ-1 | 4 | 57.9 | 17.6 |
| 367.5466 | 575.2936 | 15.7 | e3.NifQ.LWLLYHAGK.+3y5.light | NifQ-1 | 4 | 57.9 | 17.6 |
| 370.218 | 696.3919 | 15.7 | e3.NifQ.LWLLYHAGK.+3y6.heavy | NifQ-1 | 4 | 57.9 | 17.6 |
| 367.5466 | 688.3777 | 15.7 | e3.NifQ.LWLLYHAGK.+3y6.light | NifQ-1 | 4 | 57.9 | 17.6 |
| 335.6827 | 369.212 | 5.5 | e3.NifQ.MGLSPR.+2y3.heavy | NifQ-2 | 4 | 55.2 | 20.8 |
| 330.6785 | 359.2037 | 5.5 | e3.NifQ.MGLSPR.+2y3.light | NifQ-2 | 4 | 55.2 | 20.8 |
| 335.6827 | 482.2961 | 5.5 | e3.NifQ.MGLSPR.+2y4.heavy | NifQ-2 | 4 | 55.2 | 20.8 |
| 330.6785 | 472.2878 | 5.5 | e3.NifQ.MGLSPR.+2y4.light | NifQ-2 | 4 | 55.2 | 20.8 |
| **335.6827** | **539.3175** | **5.5** | **e3.NifQ.MGLSPR.+2y5.heavy** | **NifQ-2** | **4** | **55.2** | **20.8** |
| **330.6785** | **529.3093** | **5.5** | **e3.NifQ.MGLSPR.+2y5.light** | **NifQ-2** | **4** | **55.2** | **20.8** |
| 386.7206 | 446.2386 | 7.8 | e3.NifU.AAIANFR.+2y3.heavy | NifU-1 | 2 | 58.9 | 22.6 |
| 381.7165 | 436.2303 | 7.8 | e3.NifU.AAIANFR.+2y3.light | NifU-1 | 2 | 58.9 | 22.6 |
| 386.7206 | 517.2757 | 7.8 | e3.NifU.AAIANFR.+2y4.heavy | NifU-1 | 2 | 58.9 | 22.6 |
| 381.7165 | 507.2674 | 7.8 | e3.NifU.AAIANFR.+2y4.light | NifU-1 | 2 | 58.9 | 22.6 |
| 386.7206 | 630.3597 | 7.8 | e3.NifU.AAIANFR.+2y5.heavy | NifU-1 | 2 | 58.9 | 22.6 |
| 381.7165 | 620.3515 | 7.8 | e3.NifU.AAIANFR.+2y5.light | NifU-1 | 2 | 58.9 | 22.6 |
| 471.7264 | 543.2913 | 6.9 | e3.NifU.DHFFNPR.+2y4.heavy | NifU-2 | 4 | 65.1 | 25.7 |
| 466.7223 | 533.2831 | 6.9 | e3.NifU.DHFFNPR.+2y4.light | NifU-2 | 4 | 65.1 | 25.7 |
| 471.7264 | 690.3597 | 6.9 | e3.NifU.DHFFNPR.+2y5.heavy | NifU-2 | 4 | 65.1 | 25.7 |
| 466.7223 | 680.3515 | 6.9 | e3.NifU.DHFFNPR.+2y5.light | NifU-2 | 4 | 65.1 | 25.7 |
| 471.7264 | 827.4187 | 6.9 | e3.NifU.DHFFNPR.+2y6.heavy | NifU-2 | 4 | 65.1 | 25.7 |
| 314.82 | 396.2229 | 6.9 | e3.NifU.DHFFNPR.+3y3.heavy | NifU-2 | 4 | 53.8 | 14.5 |
| 311.484 | 386.2146 | 6.9 | e3.NifU.DHFFNPR.+3y3.light | NifU-2 | 4 | 53.8 | 14.5 |
| 314.82 | 543.2913 | 6.9 | e3.NifU.DHFFNPR.+3y4.heavy | NifU-2 | 4 | 53.8 | 14.5 |
| 311.484 | 533.2831 | 6.9 | e3.NifU.DHFFNPR.+3y4.light | NifU-2 | 4 | 53.8 | 14.5 |
| 314.82 | 690.3597 | 6.9 | e3.NifU.DHFFNPR.+3y5.heavy | NifU-2 | 4 | 53.8 | 14.5 |
| 311.484 | 680.3515 | 6.9 | e3.NifU.DHFFNPR.+3y5.light | NifU-2 | 4 | 53.8 | 14.5 |
| 371.7077 | 399.259 | 3.57 | e3.NifV.ASGQTLR.+2y3.heavy | NifV-3 | 4 | 57.8 | 22.1 |
| 366.7036 | 389.2507 | 3.57 | e3.NifV.ASGQTLR.+2y3.light | NifV-3 | 4 | 57.8 | 22.1 |
| 371.7077 | 527.3175 | 3.57 | e3.NifV.ASGQTLR.+2y4.heavy | NifV-3 | 4 | 57.8 | 22.1 |
| 366.7036 | 517.3093 | 3.57 | e3.NifV.ASGQTLR.+2y4.light | NifV-3 | 4 | 57.8 | 22.1 |
| 371.7077 | 584.339 | 3.57 | e3.NifV.ASGQTLR.+2y5.heavy | NifV-3 | 4 | 57.8 | 22.1 |
| 366.7036 | 574.3307 | 3.57 | e3.NifV.ASGQTLR.+2y5.light | NifV-3 | 4 | 57.8 | 22.1 |
| 401.7023 | 455.2492 | 7.9 | e3.NifV.FAENWK.+2y3.heavy | NifV-2 | 4 | 60.1 | 23.2 |
| 397.6952 | 447.235 | 7.9 | e3.NifV.FAENWK.+2y3.light | NifV-2 | 4 | 60.1 | 23.2 |
| 401.7023 | 584.2918 | 7.9 | e3.NifV.FAENWK.+2y4.heavy | NifV-2 | 4 | 60.1 | 23.2 |
| 397.6952 | 576.2776 | 7.9 | e3.NifV.FAENWK.+2y4.light | NifV-2 | 4 | 60.1 | 23.2 |
| 401.7023 | 655.329 | 7.9 | e3.NifV.FAENWK.+2y5.heavy | NifV-2 | 4 | 60.1 | 23.2 |
| 397.6952 | 647.3148 | 7.9 | e3.NifV.FAENWK.+2y5.light | NifV-2 | 4 | 60.1 | 23.2 |
| 377.7077 | 384.2229 | 9.6 | e3.NifV.VAEAAQR.+2y3.heavy | NifV-1 | 4 | 58.3 | 22.3 |
| 372.7036 | 374.2146 | 9.6 | e3.NifV.VAEAAQR.+2y3.light | NifV-1 | 4 | 58.3 | 22.3 |

| | | | | | | | |
|---|---|---|---|---|---|---|---|
| 377.7077 | 455.26 | 9.6 | e3.NifV.VAEAAQR.+2y4.heavy | NifV-1 | 4 | 58.3 | 22.3 |
| 372.7036 | 445.2518 | 9.6 | e3.NifV.VAEAAQR.+2y4.light | NifV-1 | 4 | 58.3 | 22.3 |
| 377.7077 | 584.3026 | 9.6 | e3.NifV.VAEAAQR.+2y5.heavy | NifV-1 | 4 | 58.3 | 22.3 |
| 372.7036 | 574.2944 | 9.6 | e3.NifV.VAEAAQR.+2y5.light | NifV-1 | 4 | 58.3 | 22.3 |
| **468.7605** | **540.3128** | **6.4** | **e3.NifW.AEVPLQNR.+2y4.heavy** | **NifW-1** | **2** | **64.9** | **25.6** |
| **463.7563** | **530.3045** | **6.4** | **e3.NifW.AEVPLQNR.+2y4.light** | **NifW-1** | **2** | **64.9** | **25.6** |
| **468.7605** | **637.3656** | **6.4** | **e3.NifW.AEVPLQNR.+2y5.heavy** | **NifW-1** | **2** | **64.9** | **25.6** |
| **463.7563** | **627.3573** | **6.4** | **e3.NifW.AEVPLQNR.+2y5.light** | **NifW-1** | **2** | **64.9** | **25.6** |
| **468.7605** | **736.434** | **6.4** | **e3.NifW.AEVPLQNR.+2y6.heavy** | **NifW-1** | **2** | **64.9** | **25.6** |
| **463.7563** | **726.4257** | **6.4** | **e3.NifW.AEVPLQNR.+2y6.light** | **NifW-1** | **2** | **64.9** | **25.6** |
| 418.7545 | 482.3325 | 15.9 | e3.NifW.APWLLAR.+2y4.heavy | NifW-2 | 4 | 61.3 | 23.8 |
| 413.7503 | 472.3242 | 15.9 | e3.NifW.APWLLAR.+2y4.light | NifW-2 | 4 | 61.3 | 23.8 |
| 418.7545 | 668.4118 | 15.9 | e3.NifW.APWLLAR.+2y5.heavy | NifW-2 | 4 | 61.3 | 23.8 |
| 413.7503 | 658.4035 | 15.9 | e3.NifW.APWLLAR.+2y5.light | NifW-2 | 4 | 61.3 | 23.8 |
| 418.7545 | 765.4645 | 15.9 | e3.NifW.APWLLAR.+2y6.heavy | NifW-2 | 4 | 61.3 | 23.8 |
| 413.7503 | 755.4563 | 15.9 | e3.NifW.APWLLAR.+2y6.light | NifW-2 | 4 | 61.3 | 23.8 |
| 367.197 | 431.264 | 3.8 | e3.NifZ.GSTGFVR.+2y3.heavy | NifZ-1 | 4 | 57.5 | 21.9 |
| 362.1928 | 421.2558 | 3.8 | e3.NifZ.GSTGFVR.+2y3.light | NifZ-1 | 4 | 57.5 | 21.9 |
| 367.197 | 488.2855 | 3.8 | e3.NifZ.GSTGFVR.+2y4.heavy | NifZ-1 | 4 | 57.5 | 21.9 |
| 362.1928 | 478.2772 | 3.8 | e3.NifZ.GSTGFVR.+2y4.light | NifZ-1 | 4 | 57.5 | 21.9 |
| **367.197** | **589.3332** | **3.8** | **e3.NifZ.GSTGFVR.+2y5.heavy** | **NifZ-1** | **4** | **57.5** | **21.9** |
| **362.1928** | **579.3249** | **3.8** | **e3.NifZ.GSTGFVR.+2y5.light** | **NifZ-1** | **4** | **57.5** | **21.9** |
| 674.3971 | 1248.719 | 20.4 | e3.NifZ.VPVQAIALIEER.+2y11.heavy | NifZ-2 | 4 | 79.9 | 33 |
| 669.393 | 1238.71 | 20.4 | e3.NifZ.VPVQAIALIEER.+2y11.light | NifZ-2 | 4 | 79.9 | 33 |
| 674.3971 | 740.4176 | 20.4 | e3.NifZ.VPVQAIALIEER.+2y6.heavy | NifZ-2 | 4 | 79.9 | 33 |
| 669.393 | 730.4094 | 20.4 | e3.NifZ.VPVQAIALIEER.+2y6.light | NifZ-2 | 4 | 79.9 | 33 |
| 674.3971 | 853.5017 | 20.4 | e3.NifZ.VPVQAIALIEER.+2y7.heavy | NifZ-2 | 4 | 79.9 | 33 |
| 669.393 | 843.4934 | 20.4 | e3.NifZ.VPVQAIALIEER.+2y7.light | NifZ-2 | 4 | 79.9 | 33 |
| 674.3971 | 924.5388 | 20.4 | e3.NifZ.VPVQAIALIEER.+2y8.heavy | NifZ-2 | 4 | 79.9 | 33 |
| 669.393 | 914.5306 | 20.4 | e3.NifZ.VPVQAIALIEER.+2y8.light | NifZ-2 | 4 | 79.9 | 33 |
| **465.2809** | **632.3494** | **11.3** | **e4.NifE.ALLYTGGVK.+2y6.heavy** | **NifE-1** | **4** | **64.7** | **25.5** |
| **461.2738** | **624.3352** | **11.3** | **e4.NifE.ALLYTGGVK.+2y6.light** | **NifE-1** | **4** | **64.7** | **25.5** |
| 465.2809 | 745.4334 | 11.3 | e4.NifE.ALLYTGGVK.+2y7.heavy | NifE-1 | 4 | 64.7 | 25.5 |
| 461.2738 | 737.4192 | 11.3 | e4.NifE.ALLYTGGVK.+2y7.light | NifE-1 | 4 | 64.7 | 25.5 |
| 465.2809 | 858.5175 | 11.3 | e4.NifE.ALLYTGGVK.+2y8.heavy | NifE-1 | 4 | 64.7 | 25.5 |
| 461.2738 | 850.5033 | 11.3 | e4.NifE.ALLYTGGVK.+2y8.light | NifE-1 | 4 | 64.7 | 25.5 |
| **627.8395** | **669.3554** | **20.3** | **e4.NifE.LPFLDINQER.+2y5.heavy** | **NifE-3** | **4** | **76.5** | **31.3** |
| **622.8353** | **659.3471** | **20.3** | **e4.NifE.LPFLDINQER.+2y5.light** | **NifE-3** | **4** | **76.5** | **31.3** |
| **627.8395** | **784.3823** | **20.3** | **e4.NifE.LPFLDINQER.+2y6.heavy** | **NifE-3** | **4** | **76.5** | **31.3** |
| **622.8353** | **774.3741** | **20.3** | **e4.NifE.LPFLDINQER.+2y6.light** | **NifE-3** | **4** | **76.5** | **31.3** |
| **627.8395** | **897.4664** | **20.3** | **e4.NifE.LPFLDINQER.+2y7.heavy** | **NifE-3** | **4** | **76.5** | **31.3** |
| **622.8353** | **887.4581** | **20.3** | **e4.NifE.LPFLDINQER.+2y7.light** | **NifE-3** | **4** | **76.5** | **31.3** |
| **627.8395** | **1141.588** | **20.3** | **e4.NifE.LPFLDINQER.+2y9.heavy** | **NifE-3** | **4** | **76.5** | **31.3** |
| **622.8353** | **1131.579** | **20.3** | **e4.NifE.LPFLDINQER.+2y9.light** | **NifE-3** | **4** | **76.5** | **31.3** |
| **494.7887** | **546.3274** | **18.5** | **e4.NifE.TLLDVVYR.+2y4.heavy** | **NifE-2** | **1** | **66.8** | **26.5** |
| **489.7846** | **536.3191** | **18.5** | **e4.NifE.TLLDVVYR.+2y4.light** | **NifE-2** | **1** | **66.8** | **26.5** |
| **494.7887** | **661.3543** | **18.5** | **e4.NifE.TLLDVVYR.+2y5.heavy** | **NifE-2** | **1** | **66.8** | **26.5** |
| **489.7846** | **651.3461** | **18.5** | **e4.NifE.TLLDVVYR.+2y5.light** | **NifE-2** | **1** | **66.8** | **26.5** |

| | | | | | | | |
|---|---|---|---|---|---|---|---|
| **494.7887** | **774.4384** | **18.5** | **e4.NifE.TLLDVVYR.+2y6.heavy** | **NifE-2** | **1** | **66.8** | **26.5** |
| **489.7846** | **764.4301** | **18.5** | **e4.NifE.TLLDVVYR.+2y6.light** | **NifE-2** | **1** | **66.8** | **26.5** |
| **499.7683** | **594.3597** | **13.2** | **e4.NifM.DAFAPLAQR.+2y5.heavy** | **NifM-1** | **1** | **67.2** | **26.7** |
| **494.7642** | **584.3515** | **13.2** | **e4.NifM.DAFAPLAQR.+2y5.light** | **NifM-1** | **1** | **67.2** | **26.7** |
| 499.7683 | 665.3969 | 13.2 | e4.NifM.DAFAPLAQR.+2y6.heavy | NifM-1 | 1 | 67.2 | 26.7 |
| 494.7642 | 655.3886 | 13.2 | e4.NifM.DAFAPLAQR.+2y6.light | NifM-1 | 1 | 67.2 | 26.7 |
| 499.7683 | 812.4653 | 13.2 | e4.NifM.DAFAPLAQR.+2y7.heavy | NifM-1 | 1 | 67.2 | 26.7 |
| 494.7642 | 802.457 | 13.2 | e4.NifM.DAFAPLAQR.+2y7.light | NifM-1 | 1 | 67.2 | 26.7 |
| **564.818** | **702.3809** | **21.7** | **e4.NifM.LELAFADIAR.+2y6.heavy** | **NifM-2** | **2** | **71.9** | **29** |
| **559.8139** | **692.3726** | **21.7** | **e4.NifM.LELAFADIAR.+2y6.light** | **NifM-2** | **2** | **71.9** | **29** |
| 564.818 | 773.418 | 21.7 | e4.NifM.LELAFADIAR.+2y7.heavy | NifM-2 | 2 | 71.9 | 29 |
| 559.8139 | 763.4097 | 21.7 | e4.NifM.LELAFADIAR.+2y7.light | NifM-2 | 2 | 71.9 | 29 |
| **564.818** | **886.502** | **21.7** | **e4.NifM.LELAFADIAR.+2y8.heavy** | **NifM-2** | **2** | **71.9** | **29** |
| **559.8139** | **876.4938** | **21.7** | **e4.NifM.LELAFADIAR.+2y8.light** | **NifM-2** | **2** | **71.9** | **29** |
| **691.3893** | **494.3325** | **26.9** | **e4.NifN.DLAEQFALPLVR.+2y4.heavy** | **NifN-2** | **1** | **81.2** | **33.6** |
| **686.3852** | **484.3242** | **26.9** | **e4.NifN.DLAEQFALPLVR.+2y4.light** | **NifN-2** | **1** | **81.2** | **33.6** |
| **691.3893** | **825.5221** | **26.9** | **e4.NifN.DLAEQFALPLVR.+2y7.heavy** | **NifN-2** | **1** | **81.2** | **33.6** |
| **686.3852** | **815.5138** | **26.9** | **e4.NifN.DLAEQFALPLVR.+2y7.light** | **NifN-2** | **1** | **81.2** | **33.6** |
| **691.3893** | **953.5806** | **26.9** | **e4.NifN.DLAEQFALPLVR.+2y8.heavy** | **NifN-2** | **1** | **81.2** | **33.6** |
| **686.3852** | **943.5724** | **26.9** | **e4.NifN.DLAEQFALPLVR.+2y8.light** | **NifN-2** | **1** | **81.2** | **33.6** |
| **657.8682** | **709.4595** | **31** | **e4.NifN.DTLFELANLIR.+2y6.heavy** | **NifN-3** | **1** | **78.7** | **32.4** |
| **652.8641** | **699.4512** | **31** | **e4.NifN.DTLFELANLIR.+2y6.light** | **NifN-3** | **1** | **78.7** | **32.4** |
| **657.8682** | **838.502** | **31** | **e4.NifN.DTLFELANLIR.+2y7.heavy** | **NifN-3** | **1** | **78.7** | **32.4** |
| **652.8641** | **828.4938** | **31** | **e4.NifN.DTLFELANLIR.+2y7.light** | **NifN-3** | **1** | **78.7** | **32.4** |
| **657.8682** | **985.5705** | **31** | **e4.NifN.DTLFELANLIR.+2y8.heavy** | **NifN-3** | **1** | **78.7** | **32.4** |
| **652.8641** | **975.5622** | **31** | **e4.NifN.DTLFELANLIR.+2y8.light** | **NifN-3** | **1** | **78.7** | **32.4** |
| 376.2205 | 413.2382 | 5.4 | e4.NifN.QLPVER.+2y3.heavy | NifN-1 | 1 | 58.2 | 22.2 |
| 371.2163 | 403.23 | 5.4 | e4.NifN.QLPVER.+2y3.light | NifN-1 | 1 | 58.2 | 22.2 |
| **376.2205** | **510.291** | **5.4** | **e4.NifN.QLPVER.+2y4.heavy** | **NifN-1** | **1** | **58.2** | **22.2** |
| **371.2163** | **500.2827** | **5.4** | **e4.NifN.QLPVER.+2y4.light** | **NifN-1** | **1** | **58.2** | **22.2** |
| 376.2205 | 623.3751 | 5.4 | e4.NifN.QLPVER.+2y5.heavy | NifN-1 | 1 | 58.2 | 22.2 |
| 371.2163 | 613.3668 | 5.4 | e4.NifN.QLPVER.+2y5.light | NifN-1 | 1 | 58.2 | 22.2 |
| 312.6888 | 369.212 | 2.1 | e4.NifS.AALSPR.+2y3.heavy | NifS-1 | 1 | 53.5 | 19.9 |
| 307.6847 | 359.2037 | 2.1 | e4.NifS.AALSPR.+2y3.light | NifS-1 | 1 | 53.5 | 19.9 |
| 312.6888 | 482.2961 | 2.1 | e4.NifS.AALSPR.+2y4.heavy | NifS-1 | 1 | 53.5 | 19.9 |
| 307.6847 | 472.2878 | 2.1 | e4.NifS.AALSPR.+2y4.light | NifS-1 | 1 | 53.5 | 19.9 |
| 312.6888 | 553.3332 | 2.1 | e4.NifS.AALSPR.+2y5.heavy | NifS-1 | 1 | 53.5 | 19.9 |
| 307.6847 | 543.3249 | 2.1 | e4.NifS.AALSPR.+2y5.light | NifS-1 | 1 | 53.5 | 19.9 |
| **482.7943** | **570.3234** | **9.7** | **e4.NifS.IPIAVGQTR.+2y5.heavy** | **NifS-2** | **1** | **65.9** | **26.1** |
| **477.7902** | **560.3151** | **9.7** | **e4.NifS.IPIAVGQTR.+2y5.light** | **NifS-2** | **1** | **65.9** | **26.1** |
| **482.7943** | **641.3605** | **9.7** | **e4.NifS.IPIAVGQTR.+2y6.heavy** | **NifS-2** | **1** | **65.9** | **26.1** |
| **477.7902** | **631.3522** | **9.7** | **e4.NifS.IPIAVGQTR.+2y6.light** | **NifS-2** | **1** | **65.9** | **26.1** |
| **482.7943** | **754.4445** | **9.7** | **e4.NifS.IPIAVGQTR.+2y7.heavy** | **NifS-2** | **1** | **65.9** | **26.1** |
| **477.7902** | **744.4363** | **9.7** | **e4.NifS.IPIAVGQTR.+2y7.light** | **NifS-2** | **1** | **65.9** | **26.1** |
| **482.7943** | **851.4973** | **9.7** | **e4.NifS.IPIAVGQTR.+2y8.heavy** | **NifS-2** | **1** | **65.9** | **26.1** |
| **477.7902** | **841.489** | **9.7** | **e4.NifS.IPIAVGQTR.+2y8.light** | **NifS-2** | **1** | **65.9** | **26.1** |
| 462.7605 | 468.3168 | 6.5 | e4.NifD.ANAEAVIAR.+2y4.heavy | NifD-4 | 1 | 64.5 | 25.3 |
| 457.7563 | 458.3085 | 6.5 | e4.NifD.ANAEAVIAR.+2y4.light | NifD-4 | 1 | 64.5 | 25.3 |

| | | | | | | | |
|---|---|---|---|---|---|---|---|
| **462.7605** | **539.3539** | **6.5** | **e4.NifD.ANAEAVIAR.+2y5.heavy** | **NifD-4** | **1** | **64.5** | **25.3** |
| **457.7563** | **529.3457** | **6.5** | **e4.NifD.ANAEAVIAR.+2y5.light** | **NifD-4** | **1** | **64.5** | **25.3** |
| 462.7605 | 668.3965 | 6.5 | e4.NifD.ANAEAVIAR.+2y6.heavy | NifD-4 | 1 | 64.5 | 25.3 |
| 457.7563 | 658.3883 | 6.5 | e4.NifD.ANAEAVIAR.+2y6.light | NifD-4 | 1 | 64.5 | 25.3 |
| 546.7944 | 573.3486 | 16.7 | e4.NifT.GADLYAYIAK.+2y5.heavy | NifT-2 | 1 | 70.7 | 28.4 |
| 542.7873 | 565.3344 | 16.7 | e4.NifT.GADLYAYIAK.+2y5.light | NifT-2 | 1 | 70.7 | 28.4 |
| 546.7944 | 736.412 | 16.7 | e4.NifT.GADLYAYIAK.+2y6.heavy | NifT-2 | 1 | 70.7 | 28.4 |
| 542.7873 | 728.3978 | 16.7 | e4.NifT.GADLYAYIAK.+2y6.light | NifT-2 | 1 | 70.7 | 28.4 |
| **546.7944** | **849.496** | **16.7** | **e4.NifT.GADLYAYIAK.+2y7.heavy** | **NifT-2** | **1** | **70.7** | **28.4** |
| **542.7873** | **841.4818** | **16.7** | **e4.NifT.GADLYAYIAK.+2y7.light** | **NifT-2** | **1** | **70.7** | **28.4** |
| 556.7898 | 628.3288 | 14.1 | e4.NifT.WGGAISLEGGR.+2y6.heavy | NifT-1 | 2 | 71.3 | 28.7 |
| 551.7856 | 618.3206 | 14.1 | e4.NifT.WGGAISLEGGR.+2y6.light | NifT-1 | 2 | 71.3 | 28.7 |
| **556.7898** | **741.4129** | **14.1** | **e4.NifT.WGGAISLEGGR.+2y7.heavy** | **NifT-1** | **2** | **71.3** | **28.7** |
| **551.7856** | **731.4046** | **14.1** | **e4.NifT.WGGAISLEGGR.+2y7.light** | **NifT-1** | **2** | **71.3** | **28.7** |
| 556.7898 | 812.45 | 14.1 | e4.NifT.WGGAISLEGGR.+2y8.heavy | NifT-1 | 2 | 71.3 | 28.7 |
| 551.7856 | 802.4417 | 14.1 | e4.NifT.WGGAISLEGGR.+2y8.light | NifT-1 | 2 | 71.3 | 28.7 |
| **393.2542** | **474.2802** | **10** | **e4.NifX.LVVYGVK.+2y4.heavy** | **NifX-1** | **4** | **59.5** | **22.9** |
| **389.2471** | **466.266** | **10** | **e4.NifX.LVVYGVK.+2y4.light** | **NifX-1** | **4** | **59.5** | **22.9** |
| **393.2542** | **573.3486** | **10** | **e4.NifX.LVVYGVK.+2y5.heavy** | **NifX-1** | **4** | **59.5** | **22.9** |
| **389.2471** | **565.3344** | **10** | **e4.NifX.LVVYGVK.+2y5.light** | **NifX-1** | **4** | **59.5** | **22.9** |
| **393.2542** | **672.417** | **10** | **e4.NifX.LVVYGVK.+2y6.heavy** | **NifX-1** | **4** | **59.5** | **22.9** |
| **389.2471** | **664.4028** | **10** | **e4.NifX.LVVYGVK.+2y6.light** | **NifX-1** | **4** | **59.5** | **22.9** |
| 513.2499 | 550.2495 | 8.5 | e4.NifX.VAFASSDYR.+2y4.heavy | NifX-2 | 4 | 68.2 | 27.2 |
| 508.2458 | 540.2413 | 8.5 | e4.NifX.VAFASSDYR.+2y4.light | NifX-2 | 4 | 68.2 | 27.2 |
| 513.2499 | 637.2815 | 8.5 | e4.NifX.VAFASSDYR.+2y5.heavy | NifX-2 | 4 | 68.2 | 27.2 |
| 508.2458 | 627.2733 | 8.5 | e4.NifX.VAFASSDYR.+2y5.light | NifX-2 | 4 | 68.2 | 27.2 |
| **513.2499** | **708.3187** | **8.5** | **e4.NifX.VAFASSDYR.+2y6.heavy** | **NifX-2** | **4** | **68.2** | **27.2** |
| **508.2458** | **698.3104** | **8.5** | **e4.NifX.VAFASSDYR.+2y6.light** | **NifX-2** | **4** | **68.2** | **27.2** |
| **441.7132** | **556.2965** | **6.4** | **e4.NifY.DNPLEER.+2y4.heavy** | **NifY-1** | **2** | **63** | **24.6** |
| **436.7091** | **546.2882** | **6.4** | **e4.NifY.DNPLEER.+2y4.light** | **NifY-1** | **2** | **63** | **24.6** |
| **441.7132** | **653.3492** | **6.4** | **e4.NifY.DNPLEER.+2y5.heavy** | **NifY-1** | **2** | **63** | **24.6** |
| **436.7091** | **643.341** | **6.4** | **e4.NifY.DNPLEER.+2y5.light** | **NifY-1** | **2** | **63** | **24.6** |
| 441.7132 | 767.3922 | 6.4 | e4.NifY.DNPLEER.+2y6.heavy | NifY-1 | 2 | 63 | 24.6 |
| 436.7091 | 757.3839 | 6.4 | e4.NifY.DNPLEER.+2y6.light | NifY-1 | 2 | 63 | 24.6 |
| 416.7622 | 525.3275 | 15.4 | e4.NifY.LPPWLAK.+2y4.heavy | NifY-2 | 2 | 61.2 | 23.7 |
| 412.7551 | 517.3133 | 15.4 | e4.NifY.LPPWLAK.+2y4.light | NifY-2 | 2 | 61.2 | 23.7 |
| **416.7622** | **622.3803** | **15.4** | **e4.NifY.LPPWLAK.+2y5.heavy** | **NifY-2** | **2** | **61.2** | **23.7** |
| **412.7551** | **614.3661** | **15.4** | **e4.NifY.LPPWLAK.+2y5.light** | **NifY-2** | **2** | **61.2** | **23.7** |
| **416.7622** | **719.433** | **15.4** | **e4.NifY.LPPWLAK.+2y6.heavy** | **NifY-2** | **2** | **61.2** | **23.7** |
| **412.7551** | **711.4188** | **15.4** | **e4.NifY.LPPWLAK.+2y6.light** | **NifY-2** | **2** | **61.2** | **23.7** |
| 511.8204 | 609.4061 | 19 | e45.BSA.QTALVELLK.+2y5.heavy | BSA-3 | 0.5 | 68.1 | 27.1 |
| 507.8133 | 601.3919 | 19 | e45.BSA.QTALVELLK.+2y5.light | BSA-3 | 0.5 | 68.1 | 27.1 |
| 511.8204 | 722.4902 | 19 | e45.BSA.QTALVELLK.+2y6.heavy | BSA-3 | 0.5 | 68.1 | 27.1 |
| 507.8133 | 714.476 | 19 | e45.BSA.QTALVELLK.+2y6.light | BSA-3 | 0.5 | 68.1 | 27.1 |
| 511.8204 | 793.5273 | 19 | e45.BSA.QTALVELLK.+2y7.heavy | BSA-3 | 0.5 | 68.1 | 27.1 |
| 507.8133 | 785.5131 | 19 | e45.BSA.QTALVELLK.+2y7.light | BSA-3 | 0.5 | 68.1 | 27.1 |
| **491.2838** | **595.4165** | **11.6** | **e5.NifB.EAGEILIAR.+2y5.heavy** | **NifB-1** | **1** | **66.6** | **26.4** |
| **486.2796** | **585.4083** | **11.6** | **e5.NifB.EAGEILIAR.+2y5.light** | **NifB-1** | **1** | **66.6** | **26.4** |

| | | | | | | | |
|---|---|---|---|---|---|---|---|
| **491.2838** | **724.4591** | **11.6** | **e5.NifB.EAGEILIAR.+2y6.heavy** | **NifB-1** | **1** | **66.6** | **26.4** |
| **486.2796** | **714.4509** | **11.6** | **e5.NifB.EAGEILIAR.+2y6.light** | **NifB-1** | **1** | **66.6** | **26.4** |
| **491.2838** | **781.4806** | **11.6** | **e5.NifB.EAGEILIAR.+2y7.heavy** | **NifB-1** | **1** | **66.6** | **26.4** |
| **486.2796** | **771.4723** | **11.6** | **e5.NifB.EAGEILIAR.+2y7.light** | **NifB-1** | **1** | **66.6** | **26.4** |
| 377.7385 | 411.2953 | 11.9 | e5.NifB.TLELIR.+2y3.heavy | NifB-2 | 1 | 58.3 | 22.3 |
| 372.7343 | 401.2871 | 11.9 | e5.NifB.TLELIR.+2y3.light | NifB-2 | 1 | 58.3 | 22.3 |
| **377.7385** | **540.3379** | **11.9** | **e5.NifB.TLELIR.+2y4.heavy** | **NifB-2** | **1** | **58.3** | **22.3** |
| **372.7343** | **530.3297** | **11.9** | **e5.NifB.TLELIR.+2y4.light** | **NifB-2** | **1** | **58.3** | **22.3** |
| **377.7385** | **653.422** | **11.9** | **e5.NifB.TLELIR.+2y5.heavy** | **NifB-2** | **1** | **58.3** | **22.3** |
| **372.7343** | **643.4137** | **11.9** | **e5.NifB.TLELIR.+2y5.light** | **NifB-2** | **1** | **58.3** | **22.3** |
| 563.3591 | 381.2484 | 10.5 | e5.NifD.ALDKPVIPVR.+2y3.heavy | NifD-2 | 1 | 71.5 | 28.8 |
| 554.3479 | 371.2401 | 10.5 | e5.NifD.ALDKPVIPVR.+2y3.light | NifD-2 | 1 | 71.5 | 28.8 |
| 563.3591 | 593.4009 | 10.5 | e5.NifD.ALDKPVIPVR.+2y5.heavy | NifD-2 | 1 | 71.5 | 28.8 |
| 554.3479 | 583.3926 | 10.5 | e5.NifD.ALDKPVIPVR.+2y5.light | NifD-2 | 1 | 71.5 | 28.8 |
| **563.3591** | **690.4536** | **10.5** | **e5.NifD.ALDKPVIPVR.+2y6.heavy** | **NifD-2** | **1** | **71.5** | **28.8** |
| **554.3479** | **680.4454** | **10.5** | **e5.NifD.ALDKPVIPVR.+2y6.light** | **NifD-2** | **1** | **71.5** | **28.8** |
| **563.3591** | **826.5628** | **10.5** | **e5.NifD.ALDKPVIPVR.+2y7.heavy** | **NifD-2** | **1** | **71.5** | **28.8** |
| **554.3479** | **808.5403** | **10.5** | **e5.NifD.ALDKPVIPVR.+2y7.light** | **NifD-2** | **1** | **71.5** | **28.8** |
| 614.3857 | 695.4542 | 12.4 | e5.NifD.ALKPDLIGSGIK.+2y7.heavy | NifD-3 | 1 | 75.3 | 30.7 |
| 606.3715 | 687.44 | 12.4 | e5.NifD.ALKPDLIGSGIK.+2y7.light | NifD-3 | 1 | 75.3 | 30.7 |
| 614.3857 | 810.4811 | 12.4 | e5.NifD.ALKPDLIGSGIK.+2y8.heavy | NifD-3 | 1 | 75.3 | 30.7 |
| 606.3715 | 802.4669 | 12.4 | e5.NifD.ALKPDLIGSGIK.+2y8.light | NifD-3 | 1 | 75.3 | 30.7 |
| **614.3857** | **907.5339** | **12.4** | **e5.NifD.ALKPDLIGSGIK.+2y9.heavy** | **NifD-3** | **1** | **75.3** | **30.7** |
| **606.3715** | **899.5197** | **12.4** | **e5.NifD.ALKPDLIGSGIK.+2y9.light** | **NifD-3** | **1** | **75.3** | **30.7** |
| 412.7596 | 365.2638 | 13.9 | e5.NifD.GVVFGPIK.+2y3.heavy | NifD-1 | 1 | 60.9 | 23.6 |
| 408.7525 | 357.2496 | 13.9 | e5.NifD.GVVFGPIK.+2y3.light | NifD-1 | 1 | 60.9 | 23.6 |
| **412.7596** | **422.2853** | **13.9** | **e5.NifD.GVVFGPIK.+2y4.heavy** | **NifD-1** | **1** | **60.9** | **23.6** |
| **408.7525** | **414.2711** | **13.9** | **e5.NifD.GVVFGPIK.+2y4.light** | **NifD-1** | **1** | **60.9** | **23.6** |
| **412.7596** | **569.3537** | **13.9** | **e5.NifD.GVVFGPIK.+2y5.heavy** | **NifD-1** | **1** | **60.9** | **23.6** |
| **408.7525** | **561.3395** | **13.9** | **e5.NifD.GVVFGPIK.+2y5.light** | **NifD-1** | **1** | **60.9** | **23.6** |
| **412.7596** | **668.4221** | **13.9** | **e5.NifD.GVVFGPIK.+2y6.heavy** | **NifD-1** | **1** | **60.9** | **23.6** |
| **408.7525** | **660.4079** | **13.9** | **e5.NifD.GVVFGPIK.+2y6.light** | **NifD-1** | **1** | **60.9** | **23.6** |
| **760.4088** | **722.4183** | **16.6** | **e5.NifF.QLGELADAPVNINR.+2y6.heavy** | **NifF-2** | **1** | **86.2** | **36.1** |
| **755.4046** | **712.41** | **16.6** | **e5.NifF.QLGELADAPVNINR.+2y6.light** | **NifF-2** | **1** | **86.2** | **36.1** |
| **760.4088** | **793.4554** | **16.6** | **e5.NifF.QLGELADAPVNINR.+2y7.heavy** | **NifF-2** | **1** | **86.2** | **36.1** |
| **755.4046** | **783.4472** | **16.6** | **e5.NifF.QLGELADAPVNINR.+2y7.light** | **NifF-2** | **1** | **86.2** | **36.1** |
| **760.4088** | **908.4824** | **16.6** | **e5.NifF.QLGELADAPVNINR.+2y8.heavy** | **NifF-2** | **1** | **86.2** | **36.1** |
| **755.4046** | **898.4741** | **16.6** | **e5.NifF.QLGELADAPVNINR.+2y8.light** | **NifF-2** | **1** | **86.2** | **36.1** |
| **760.4088** | **979.5195** | **16.6** | **e5.NifF.QLGELADAPVNINR.+2y9.heavy** | **NifF-2** | **1** | **86.2** | **36.1** |
| **755.4046** | **969.5112** | **16.6** | **e5.NifF.QLGELADAPVNINR.+2y9.light** | **NifF-2** | **1** | **86.2** | **36.1** |
| **593.8263** | **655.3397** | **19** | **e5.NifF.TVALFGLGDQR.+2y6.heavy** | **NifF-1** | **1** | **74** | **30.1** |
| **588.8222** | **645.3315** | **19** | **e5.NifF.TVALFGLGDQR.+2y6.light** | **NifF-1** | **1** | **74** | **30.1** |
| **593.8263** | **802.4081** | **19** | **e5.NifF.TVALFGLGDQR.+2y7.heavy** | **NifF-1** | **1** | **74** | **30.1** |
| **588.8222** | **792.3999** | **19** | **e5.NifF.TVALFGLGDQR.+2y7.light** | **NifF-1** | **1** | **74** | **30.1** |
| **593.8263** | **915.4922** | **19** | **e5.NifF.TVALFGLGDQR.+2y8.heavy** | **NifF-1** | **1** | **74** | **30.1** |
| **588.8222** | **905.4839** | **19** | **e5.NifF.TVALFGLGDQR.+2y8.light** | **NifF-1** | **1** | **74** | **30.1** |
| **377.7077** | **412.2542** | **2.9** | **e5.NifH.DNIVQR.+2y3.heavy** | **NifH-2** | **1** | **58.3** | **22.3** |
| **372.7036** | **402.2459** | **2.9** | **e5.NifH.DNIVQR.+2y3.light** | **NifH-2** | **1** | **58.3** | **22.3** |

| | | | | | | | |
|---|---|---|---|---|---|---|---|
| **377.7077** | **525.3383** | **2.9** | **e5.NifH.DNIVQR.+2y4.heavy** | **NifH-2** | **1** | **58.3** | **22.3** |
| **372.7036** | **515.33** | **2.9** | **e5.NifH.DNIVQR.+2y4.light** | **NifH-2** | **1** | **58.3** | **22.3** |
| 377.7077 | 639.3812 | 2.9 | e5.NifH.DNIVQR.+2y5.heavy | NifH-2 | 1 | 58.3 | 22.3 |
| 372.7036 | 629.3729 | 2.9 | e5.NifH.DNIVQR.+2y5.light | NifH-2 | 1 | 58.3 | 22.3 |
| **351.7412** | **363.223** | **3.4** | **e5.NifH.LILHAK.+2y3.heavy** | **NifH-1** | **1** | **56.5** | **21.4** |
| **347.7341** | **355.2088** | **3.4** | **e5.NifH.LILHAK.+2y3.light** | **NifH-1** | **1** | **56.5** | **21.4** |
| **351.7412** | **476.3071** | **3.4** | **e5.NifH.LILHAK.+2y4.heavy** | **NifH-1** | **1** | **56.5** | **21.4** |
| **347.7341** | **468.2929** | **3.4** | **e5.NifH.LILHAK.+2y4.light** | **NifH-1** | **1** | **56.5** | **21.4** |
| **351.7412** | **589.3912** | **3.4** | **e5.NifH.LILHAK.+2y5.heavy** | **NifH-1** | **1** | **56.5** | **21.4** |
| **347.7341** | **581.377** | **3.4** | **e5.NifH.LILHAK.+2y5.light** | **NifH-1** | **1** | **56.5** | **21.4** |
| **536.8049** | **617.3281** | **18.5** | **e5.NifJ.LITQLFGDR.+2y5.heavy** | **NifJ-2** | **1** | **69.9** | **28** |
| **531.8007** | **607.3198** | **18.5** | **e5.NifJ.LITQLFGDR.+2y5.light** | **NifJ-2** | **1** | **69.9** | **28** |
| **536.8049** | **745.3867** | **18.5** | **e5.NifJ.LITQLFGDR.+2y6.heavy** | **NifJ-2** | **1** | **69.9** | **28** |
| **531.8007** | **735.3784** | **18.5** | **e5.NifJ.LITQLFGDR.+2y6.light** | **NifJ-2** | **1** | **69.9** | **28** |
| **536.8049** | **846.4344** | **18.5** | **e5.NifJ.LITQLFGDR.+2y7.heavy** | **NifJ-2** | **1** | **69.9** | **28** |
| **531.8007** | **836.4261** | **18.5** | **e5.NifJ.LITQLFGDR.+2y7.light** | **NifJ-2** | **1** | **69.9** | **28** |
| 470.7656 | 381.2484 | 11.5 | e5.NifJ.NLFGQPVR.+2y3.heavy | NifJ-1 | 1 | 65.1 | 25.6 |
| 465.7614 | 371.2401 | 11.5 | e5.NifJ.NLFGQPVR.+2y3.light | NifJ-1 | 1 | 65.1 | 25.6 |
| 470.7656 | 509.307 | 11.5 | e5.NifJ.NLFGQPVR.+2y4.heavy | NifJ-1 | 1 | 65.1 | 25.6 |
| 465.7614 | 499.2987 | 11.5 | e5.NifJ.NLFGQPVR.+2y4.light | NifJ-1 | 1 | 65.1 | 25.6 |
| **470.7656** | **566.3284** | **11.5** | **e5.NifJ.NLFGQPVR.+2y5.heavy** | **NifJ-1** | **1** | **65.1** | **25.6** |
| **465.7614** | **556.3202** | **11.5** | **e5.NifJ.NLFGQPVR.+2y5.light** | **NifJ-1** | **1** | **65.1** | **25.6** |
| **470.7656** | **713.3969** | **11.5** | **e5.NifJ.NLFGQPVR.+2y6.heavy** | **NifJ-1** | **1** | **65.1** | **25.6** |
| **465.7614** | **703.3886** | **11.5** | **e5.NifJ.NLFGQPVR.+2y6.light** | **NifJ-1** | **1** | **65.1** | **25.6** |
| **477.786** | **508.3481** | **19.6** | **e5.NifK.AFEVPLIR.+2y4.heavy** | **NifK-2** | **0.5** | **65.6** | **25.9** |
| **472.7818** | **498.3398** | **19.6** | **e5.NifK.AFEVPLIR.+2y4.light** | **NifK-2** | **0.5** | **65.6** | **25.9** |
| **477.786** | **607.4165** | **19.6** | **e5.NifK.AFEVPLIR.+2y5.heavy** | **NifK-2** | **0.5** | **65.6** | **25.9** |
| **472.7818** | **597.4083** | **19.6** | **e5.NifK.AFEVPLIR.+2y5.light** | **NifK-2** | **0.5** | **65.6** | **25.9** |
| **477.786** | **736.4591** | **19.6** | **e5.NifK.AFEVPLIR.+2y6.heavy** | **NifK-2** | **0.5** | **65.6** | **25.9** |
| **472.7818** | **726.4509** | **19.6** | **e5.NifK.AFEVPLIR.+2y6.light** | **NifK-2** | **0.5** | **65.6** | **25.9** |
| **476.2655** | **323.2169** | **8** | **e5.NifK.EALTVDPAK.+2y3.heavy** | **NifK-1** | **0.5** | **65.5** | **25.9** |
| **472.2584** | **315.2027** | **8** | **e5.NifK.EALTVDPAK.+2y3.light** | **NifK-1** | **0.5** | **65.5** | **25.9** |
| **476.2655** | **537.3122** | **8** | **e5.NifK.EALTVDPAK.+2y5.heavy** | **NifK-1** | **0.5** | **65.5** | **25.9** |
| **472.2584** | **529.298** | **8** | **e5.NifK.EALTVDPAK.+2y5.light** | **NifK-1** | **0.5** | **65.5** | **25.9** |
| **476.2655** | **638.3599** | **8** | **e5.NifK.EALTVDPAK.+2y6.heavy** | **NifK-1** | **0.5** | **65.5** | **25.9** |
| **472.2584** | **630.3457** | **8** | **e5.NifK.EALTVDPAK.+2y6.light** | **NifK-1** | **0.5** | **65.5** | **25.9** |
| **476.2655** | **751.444** | **8** | **e5.NifK.EALTVDPAK.+2y7.heavy** | **NifK-1** | **0.5** | **65.5** | **25.9** |
| **472.2584** | **743.4298** | **8** | **e5.NifK.EALTVDPAK.+2y7.light** | **NifK-1** | **0.5** | **65.5** | **25.9** |
| **487.7703** | **560.3066** | **23** | **e5.NifK.LGFPLFDR.+2y4.heavy** | **NifK-3** | **0.5** | **66.3** | **26.2** |
| **482.7662** | **550.2984** | **23** | **e5.NifK.LGFPLFDR.+2y4.light** | **NifK-3** | **0.5** | **66.3** | **26.2** |
| **487.7703** | **657.3594** | **23** | **e5.NifK.LGFPLFDR.+2y5.heavy** | **NifK-3** | **0.5** | **66.3** | **26.2** |
| **482.7662** | **647.3511** | **23** | **e5.NifK.LGFPLFDR.+2y5.light** | **NifK-3** | **0.5** | **66.3** | **26.2** |
| 487.7703 | 804.4278 | 23 | e5.NifK.LGFPLFDR.+2y6.heavy | NifK-3 | 0.5 | 66.3 | 26.2 |
| 482.7662 | 794.4196 | 23 | e5.NifK.LGFPLFDR.+2y6.light | NifK-3 | 0.5 | 66.3 | 26.2 |
| 399.2466 | 484.2857 | 8.2 | eAll.BSA.LVTDLTK.+2y4.heavy | BSA-1 | 1 | 59.9 | 23.1 |
| 395.2395 | 476.2715 | 8.2 | eAll.BSA.LVTDLTK.+2y4.light | BSA-1 | 1 | 59.9 | 23.1 |
| 399.2466 | 585.3334 | 8.2 | eAll.BSA.LVTDLTK.+2y5.heavy | BSA-1 | 1 | 59.9 | 23.1 |
| 395.2395 | 577.3192 | 8.2 | eAll.BSA.LVTDLTK.+2y5.light | BSA-1 | 1 | 59.9 | 23.1 |

| | | | | | | | |
|---|---|---|---|---|---|---|---|
| 399.2466 | 684.4018 | 8.2 | eAll.BSA.LVTDLTK.+2y6.heavy | BSA-1 | 1 | 59.9 | 23.1 |
| 395.2395 | 676.3876 | 8.2 | eAll.BSA.LVTDLTK.+2y6.light | BSA-1 | 1 | 59.9 | 23.1 |

Supplementary Text 1: Calculation of counts numbers per cell

Calculations of protein count per cell were undertaken as follows:

1. Sample (light) to QconCAT (heavy) peak area ratio was determined for each transition (a)
2. (a) x final QconCAT concentration in each digestion reaction (High - 0.0007 mg/ml, Medium – 0.000124 mg/ml, Low – 0.000082 mg/ml) calculates peptide abundance in mg/ml (b)
3. $OD_{600}$ of each sample multiplied by $1.1 \times 10^9$ to give the number of *K. oxytoca* cells in 1 ml (7), and then in 30 µl (c)
4. Amount (mg) of each peptide in 30 µl was calculated from (b), before being converted to kDa (d)
5. Nif protein count in 30 µl was determined by (d)/specific Nif protein kDa, obtained from the Protein Molecular Weight calculator at Bioinformatics.org (8) (e).
6. Number of proteins per cell was determined by (e)/(c).

The same calculation was applied to the peak area corresponding to % Nif protein sample below which signal was no longer linear, in order to define quantification limit in terms of protein count per cell.

Calculation of Nif protein values per total protein involved steps 1 and 2 above. The result of step 2 was then divided by total protein as established by DC assay.

**Appendix IV: Further details on MS methods used for absolute quantification of Nif proteins**

*GC-MS*

Acetylene reduction was measured by GC-MS using an Agilent 7820 GC System with oven at 70 °C, FID at 250 °C, 6-minute run time, Nitrogen carrier gas. A 500 µl syringe with a PTFE septum was used to inject culture sample headspace samples onto a HayeSep N Column (Divinylbenzene/ethylene glycol dimethacrylate packing) (Agilent).

*MRM-MS*

Samples (typically 40 µl) were analysed by triple quadrupole HPLC-electrospray ionisation/MS-MS using a Shimadzu Prominence UPLC coupled to an Applied Biosystems Q-TRAP 6500 (AB SCIEX, Massachussetts, USA). Chromatographic separation was carried out by reverse phase chromatography on a Phenomenex Luna C18(2) 100A (3 µm, 100 by 2 mm) HPLC column, at 50 °C. A gradient system using A (5 % $CH_3CN$, 94.9 % $H_2O$, 0.1 % formic acid) and B (5 % $H_2O$, 94.9 % $CH_3CN$, 0.1 % formic acid) was used, with solvent flow rate of 250 µl/minute. A linear gradient from 0 % B to 25 % B over 30 minutes was followed by an increase to 50 % B over the next 5 minutes. B was then increased to 100 % over the next 2 minutes and held at 100 % for 5 minutes, before return to initial conditions.

The MS was operated in the positive mode using an IonDrive™ Turbo V as the ion source. Source conditions were: temperature, 500 °C; ion source gas 1, 40 psi; ion source gas 2, 60 psi; ion spray voltage, 5500 V; curtain gas, 40 psi; CAD gas setting, -2. An Enhanced Scheduled MRM method was used for the analysis, and details of the full method can be found in appendix I.

**Appendix V (next page): 15 % SDS-PAGE gels from expression and purification of Low, Medium and High QconCATs.**

(a) Resuspended cell pellets from uninduced (unind.) and IPTG-induced (ind.) *E. coli* BL 21 cell cultures during expression of stable isotope-labelled QconCATs confirm QconCAT overexpression. The gel has been stained using Coomassie. MW, PageRuler Prestained Protein Ladder (ThermoFisher Scientific). (b) Water and urea-soluble protein fractions reveal presence of QconCAT in urea-soluble fraction. MW, EZ-Ladder Fluorescent Protein Molecular Weight Markers (EZ Biolab). (c) Purification fractions from Ni-affinity purification of QconCATs show general efficient binding to the Ni resin. FT, Flow-Through; B, Binding buffer; W, Wash buffer; E, Elution Buffer; D, Post-dialysis. (d) The EZ-Ladder Fluorescent Protein Molecular Weight Markers was run with and without 7M urea to see whether urea has an effect on migration in SDS-PAGE.

Thick black or white arrows indicate the band representing each QconCAT. Approximate molecular weights (kDa) are indicated to the left (and right in the case of (d)) of each figure.

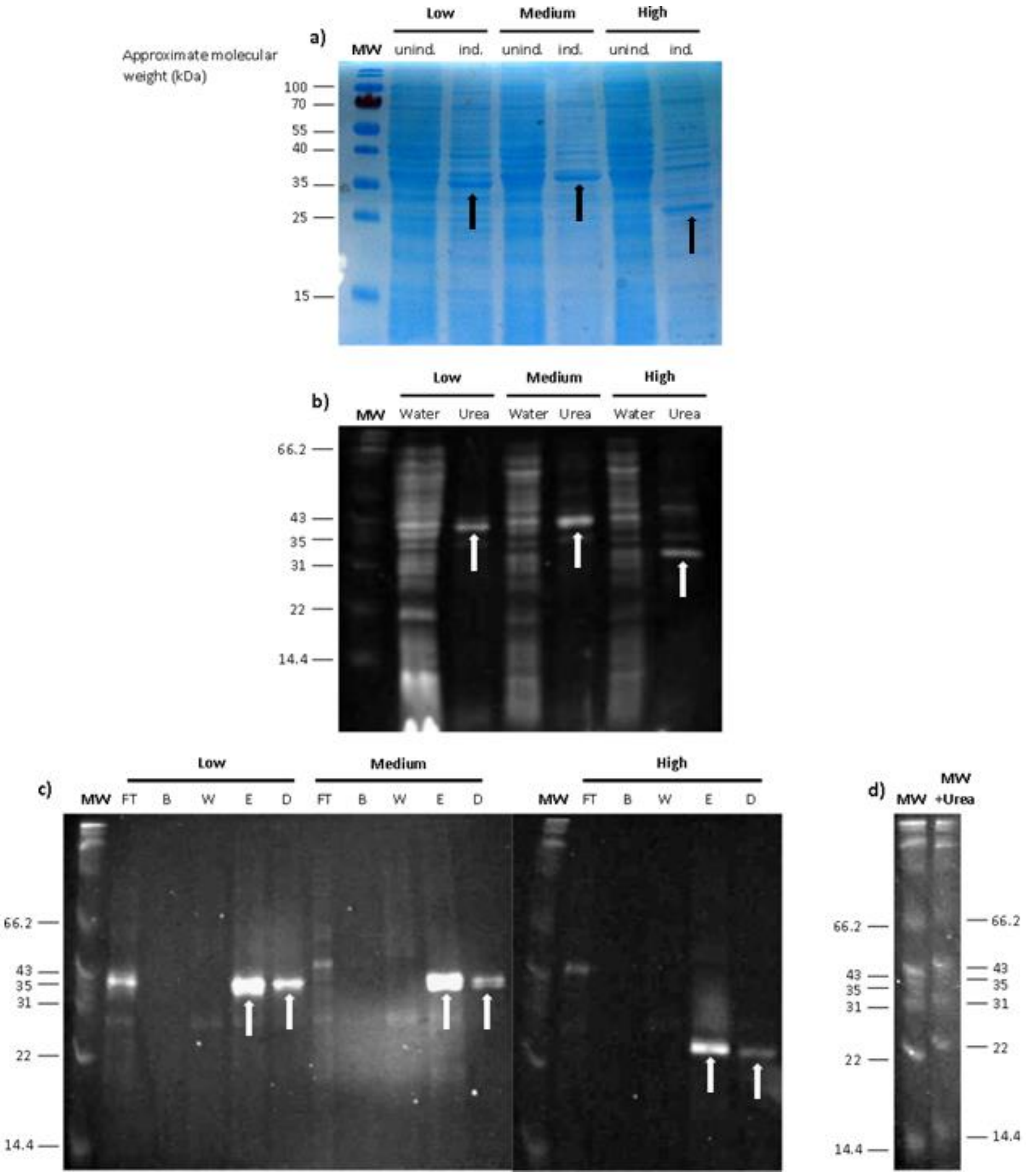
a)
Approximate molecular weight (kDa)
MW
Low
Medium
High
unind.
ind.
unind.
ind.
unind.
ind.
100
70
55
40
35
25
15
b)
MW
Low
Medium
High
Water
Urea
Water
Urea
Water
Urea
66.2
43
35
31
22
14.4
c)
Low
Medium
High
MW
FT
B
W
E
D
FT
B
W
E
D
MW
FT
B
W
E
D
66.2
43
35
31
22
14.4
d)
MW
MW +Urea
66.2
43
35
31
22
14.4
